# Genetic variability under the seedbank coalescent

**DOI:** 10.1101/017244

**Authors:** Jochen Blath, Adrián Casanova, Bjarki Eldon, Noemi Kurt, Maite Wilke-Berenguer

## Abstract

We analyse patterns of genetic variability of populations in the presence of a large seedbank with the help of a new coalescent structure called the seedbank coalescent. This ancestral process appears naturally as scaling limit of the genealogy of large populations that sustain seedbanks, if the seedbank size and individual dormancy times are of the same order as the active population. Mutations appear as Poisson processes on the active lineages, and potentially at reduced rate also on the dormant lineages. The presence of ‘dormant’ lineages leads to qualitatively altered times to the most recent common ancestor and non-classical patterns of genetic diversity. To illustrate this we provide a Wright-Fisher model with seedbank component and mutation, motivated from recent models of microbial dormancy, whose genealogy can be described by the seedbank coalescent. Based on our coalescent model, we derive recursions for the expectation and variance of the time to most recent common ancestor, number of segregating sites, pairwise differences, and singletons. Estimates (obtained by simulations) of the distributions of commonly employed distance statistics, in the presence and absence of a seedbank, are compared. The effect of a seedbank on the expected site-frequency spectrum is also investigated using simulations. Our results indicate that the presence of a large seedbank considerably alters the distribution of some distance statistics, as well as the site-frequency spectrum. Thus, one should be able to detect from genetic data the presence of a large seedbank in natural populations.

## Introduction

Many microorganisms can enter reversible dormant states of low (resp. zero) metabolic activity, for example when faced with unfavourable environmental conditions; see e.g. LENNON and JONES (2011) for a recent overview of this phenomenon. Such dormant forms may stay inactive for extended periods of time and thus create a seed bank that should significantly affect the interplay of evolutionary forces driving the genetic variability of the microbial population. In fact, in many eco-systems, the percentage of dormant cells compared to the total population size is substantial, and sometimes even dominant (for example roughly 20% in human gut, 40% in marine water, 80% in soil, cf. LENNON and JONES (2011)[Box 1, Table *a*]). This abundance of dormant forms, which can be short-lived as well as stay inactive for significant periods of time (decades or century old spores are not uncommon) thus creates a seedbank that buffers against environmental change, but potentially also against classical evolutionary forces such as genetic drift, mutation, or selection.

In this paper, we investigate the effect of large seed banks (that is, comparable to the size of the active population) on the patterns of genetic variability in populations over macroscopic timescales. In particular, we extend a recently introduced mathematical model for the ancestral relationships in a Wright-Fisherian population of size *N* with geometric seed bank age distribution (cf. BLATH *et al.* (2015b))) to accommodate different mutation rates for ‘active’ and ‘dormant’ individuals, as well as a positive death rate in the seed bank. The resulting genealogy, measured over timescales of order *N*, can then be described by a new universal coalescent structure, the ‘seed bank coalescent with mutation’, if the individual initiation and resuscitation rates between active and dormant states as well as the individual mutation rates are of order 1*/N*. Measuring times in units of *N* and mutation rates in units of 1*/N* is of course the classical scaling regime in population genetic modeling; in particular, the classical Wright-Fisher model has a genealogy that converges in precisely this setup to the usual Kingman coalescent with mutation (KINGMAN (1982a,c,b); see WAKELEY (2009) for an overview).

We will provide a precise description of these (seed bank) coalescents and corresponding population models, in part motivated by recent research in microbial dormancy JONES and LENNON (2010); LENNON and JONES (2011), in the next section below. We argue that our seed bank coalescent is universal in the sense that it is robust to the specifics of the associated population model, as long as certain basic features are captured.

Our explicit seedbank coalescent model then allows us to derive expressions for several important population genetic quantities. In particular, we provide recursions for the expectation (and variance) of the time to the most recent common ancestor (T_MRCA_), the total number of segregating sites, average pairwise differences and number of singletons in a sample (under the inifinitely-many sites model assumptions). We then use these recursions, and additional simulations based on the seedbank coalescent with mutation, to analyse Tajima’s *D* and related distance statistics in the presence of seedbanks, and also the observed site frequency spectrum.

We hope that this basic analysis triggers further research on the effect of seedbanks in population genetics, for example concerning statistical methods that allow one to infer the presence and size of seedbanks from data, to allow model selection (e.g. seedbank coalescent versus (time-changed) Kingman coalescent), and finally to estimate evolutionary parameters such as the mutation rate in dormant individuals, or the inactivation and reactivation rates between the dormant and active states.

It is important to note that our approach is different from a previously introduced mathematical seedbank model in KAJ *et al.* (2001). There, the authors consider a population of constant size *N* where each individual chooses its parent a random amount of generations in the past and copies its genetic type from there. The number of generations that separate each parent and offspring can be interpreted as the time (in generations) that the offspring stays dormant. The authors show that if the maximal time spent in the seedbank is restricted to finitely many {1, 2*, …, m*}, where *m* is fixed, then the ancestral process induced by the seedbank model converges, after the usual scaling of time by a factor *N,* to a time changed (delayed) Kingman coalescent. Thus, typical patterns of genetic diversity, in particular the normalised site frequency spectrum, will stay (qualitatively) unchanged. Of course, the point here is that the expected seedbank age distribution is not on the order of *N*, but uniformly bounded by *m*, so that for the coalescent approximation to hold one necessarily needs that *m* is small compared to *N*, which results a ‘weak’ seedbank effect. This model has been applied in TELLIER *et al.* (2011) in the analysis of seedbanks in certain species of wild tomatoes. A related model was considered in VITALIS *et al.* (2004), which shares the feature that the time spent in the seedbank is bounded by a fixed number independent of the population size. For a more detailed mathematical discussion of such models, including previous work in BLATH *et al.* (2015a), see BLATH *et al.* (2015b). The choice of the adequate coalescent model (seedbank coalescent vs. (time-changed) Kingman coalescent) will thus also be an important question for study design, and the development of corresponding model selection rules will be part of future research.

## Coalescent models and seedbanks

Before we discuss the seedbank coalescent, we briefly recall the classical Kingman coalescent for reference this will ease the comparison of the underlying assumptions of both models.

### The Kingman coalescent with mutation

The Kingman coalescent (KINGMAN, 1982a,c,b) describes the ancestral process of a large class of neutral exchangeable population models including the Wright-Fisher model (WRIGHT, 1931; FISHER, 1930), the Moran model (MORAN, 1958) and many Cannings models (CANNINGS, 1974). See e.g. WAKELEY (2009) for an overview. If we trace the ancestral lines (that is, the sequence of genetic ancestors at a locus) of a sample of size *n* backwards in time, we obtain a binary tree, in which we see pairwise coalescences of branches until the most-recent common ancestor is reached. Kingman proved that the probability law of this random tree can be describe as follows: Each pair of lineages (there are 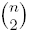 many) has the same chance to coalesce, and the successive coalescence times are exponentially distributed with parameters 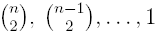 until the last remaining pair of lines has coalesced. This elegant structure allows one to easily determine the expected time to the most recent common ancestor of a sample of size *n*, which is well known to be

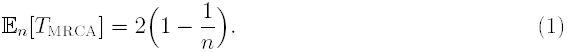

Not surprisingly, we will essentially recover (1) for the seedbank coalescent defined below if the relative seedbank size becomes small compared to the ‘active’ population size.

As usual, mutations are placed upon the resulting coalescent tree according to a Poisson-process with rate *θ/*2, for some appropriate *θ >* 0, so that the expected number of mutations of a sample of size 2 is just *θ*.

The underlying assumptions about the population for a Kingman coalescent approximation of its genealogy to be justified are simple but far-reaching, namely that the different genetic types in the population are selectively neutral (i.e. do not exhibit significant fitness differences), and that the population size of the underlying population is essentially constant in time. If the population can be described by the (haploid) Wright-Fisher model (of constant size, say *N*), then, in order to arrive at the described limiting genealogy, it is standard to measure time in units of *N*, *the coalescent time scale*, and to assume that the individual mutation rates per generation are of order *θ/*(2*N*). The exact time-scaling usually depends on the reproductive mechanism and other particularities of the underlying model (it differs already among variants of the Moran model), but the Kingman coalescent is still a universally valid limit for many a priori different population models (including e. g. all reproductive mechanisms with bounded offspring variance, dioecy, age structure, partial selfing and to some degree geographic structure), when these particularities exert their influence over time scales much shorter than the coalescent time scale, cf. e.g. WAKELEY (2013). This is also the reason, why the Kingman coalescent still appears as limiting genealogy of the ‘weak’ seedbank model of Kaj *et al.* (2001) mentioned in the introduction.

This robustness has turned the Kingman coalescent into an extremely useful tool in population genetics. In fact, it can be considered the standard null-model for neutral populations. Its success is also based on the fact that it allows a simple derivation of many population genetic quantities of interest, such as a formula for the expected number of segregating sites

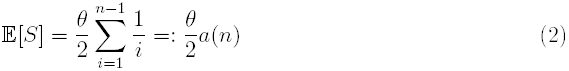

or the expected average number of pairwise differences *π* (TAJIMA, 1983), the expected values of the site-frequency spectrum, cf. Fu (1995), when one assumes the infinite-sites model of WATTEKSON (1975). This analytic tractability has allowed the construction of a sophisticated statistical machinery for the inference of evolutionary parameters. We will investigate the corresponding quantities for the seedbank coalescent below.

### The seedbank coalescent with mutation

Similar to the Kingman coalescent, the seedbank coalescent, mathematically introduced in BLATH *et al.* (2015b), describes the ancestral lines of a sample taken from a population with seedbank component. Here, we distinguish whether an ancestral line belongs to an ‘active’ or ‘dormant’ individual for any given point backward in time. The main difference to the Kingman coalescent is that as long as an ancestral line corresponds to a dormant individual (in the seedbank), it cannot coalesce with other lines, since reproduction and thus finding a common ancestor is only possible for ‘active’ individuals.

The dynamics is now easily described as follows: If there are currently *n* active and *m* dormant lineages at some point in the past, each ‘active pair’ may coalesce with the same probability, after an exponential time with rate 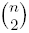, entirely similar to a classical Kingman coalescent with currently *n* lineages. However, each active line becomes dormant at a positive rate *c >* 0 (corresponding to an ancestor who emerged from the seedbank), and each dormant line resuscitates, at a rate *cK*, for some *K >* 0. The parameter *K* reflects the relative size of the seedbank compared to the active population, and will be explained below in terms of an explicit underlying population model. Since dormant lines are prevented from merging, they significantly delay the time to the most recent common ancestor. This mechanism is reminiscent of a structured coalescent with two islands (HERBOTS, 1997; NOTOHARA, 1990), where lineages may only merge if they are in the same colony. Of course, if one samples a seedbank coalescent backwards in time, one need not only specify the sample size, but actually the number of sampled individuals from the active population (say *n*), and from the dormant population (say *m*).

In this paper, we also consider mutations along the ancestral lines. As in the Kingman case we place them along the active line segments according to a Poisson process with rate *θ*_1_, and along the dormant segments at a rate *θ*_2_ *≥* 0. Depending on the concrete situation, one may want to choose *θ*_2_ = 0. To determine the mutation rate in dormant individuals will be an interesting inference question. In Figure 1, we illustrate a realisation of the seedbank coalescent with mutations: ancestral lineages residing in the seedbank are represented by dotted lines, and do not take part in coalescences.

**Figure 1:**
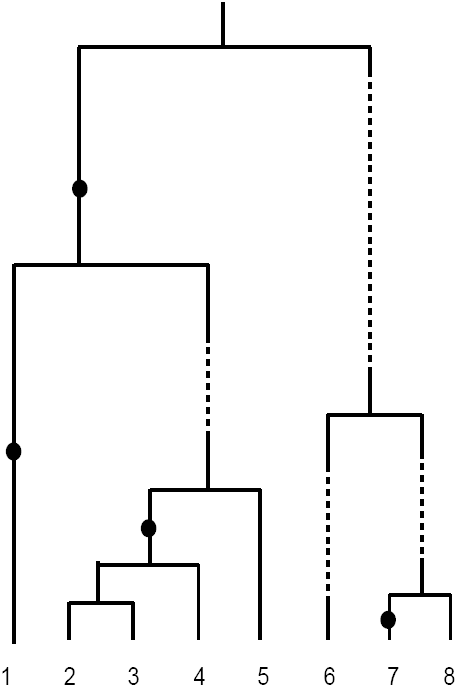
Realisation of a seedbank coalescent with all *n* = 8 sampled lineages assumed active. Mutations are, in this example, only allowed to occur on active ancestral lineages represented by solid lines; ancestral lineages residing in the seedbank are represented by dotted lines. Coalescences are only allowed among active lineages.

A formal mathematical definition of this process as partition-valued Markov chain can be found in BLATH *et al.* (2015b); it is straightforward to extend their framework to include mutations.

The parameters *c* and *K* can be understood as follows: *c* describes the proportion of individuals that enter the seedbank per (macroscopic) coalescent time-unit. It is thus the rate at which individuals become dormant. If the ratio of the size of the active population and the dormant population in the underlying population is *K*: 1 (that is, the active population is *K* times the size of the dormant population), and absolute (and thus also relative) population sizes are assumed to stay constant, then, in order for the relative amount of active and dormant individuals to stay balanced, the rate at which dormant individuals resuscitate and return to the active population is necessarily of the form *cK*, see also Figure 2. It is important to note that in this setup, the average coalescent time that an inactive individual stays dormant is of the order *N/*(*cK*). We will later also include a positive mortality rate for dormant individuals, this will lead to a reduced ‘effective’ relative seedbank 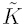.

**Figure 2:**
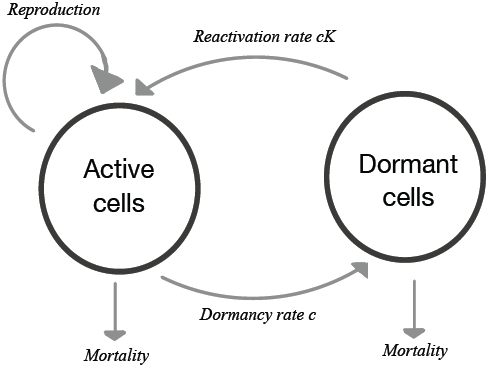
Dynamics of reversible microbial dormancy, according to JONES and LENNON (2010)

### Robustness and underlying assumptions of the seedbank coalescent

As for the Kingman coalescent, it is important to understand the underlying assumptions that make the seedbank coalescent a reasonable model for the genealogy of a population: Again, we assume the types in the population to be selectively neutral, so that there are no significant fitness differences. Further, we assume the population size *N* and the seedbank size *M* to be constant, and to be of the same order, that is there exists a *K >* 0 so that *N* = *K · M*, i.e. the ratio between active and dormant individuals is constant equal to *K*: 1. Finally, the rate at which an active individual becomes dormant should be *c* (on the macroscopic coalescent scale), so that necessarily the average time (in coalescent time units) that an individual stays dormant before being resuscitated becomes 1*/*(*cK*). If one includes a positive mortality rate in the seedbank, this will lead to a modified parameter 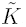, see below.

We will provide below an example of a concrete seedbank population model, the ‘Wright-Fisher model with geometric seedbank component’, including mutation and mortality in the seedbank, for which it can be proved that the seedbank coalescent with mutation governs the genealogy if the population size *N* (and thus necessarily also seedbank size *M*) gets large, and coalescent time is measured in units of the population size *N*. This is the same scaling regime as in the case of the Kingman coalescent corresponding to genealogy of the classical Wright-Fisher model.

The seedbank coalescent with mutation should be robust against small alterations – such as in the transition or reproduction mechanism, or in the population or seedbank size – of the underlying population, similar to the robustness of the Kingman coalescent. Especially if these alterations occur on time scales that are much shorter than the coalescent time scale (which is *N* for the haploid Wright-Fisher model). For example, one can still obtain this coalescent in a *Moran model* with seedbank component, as long as the seedbank is on the same order as the active population, and if the migration rates between seedbank and active population scale suitably (as well as the mutation rate) with the coalescent time scale. As mentioned above, this is an important difference to the model considered by Kaj *et al.* (2001), where the time an individual stays in the seedbank is negligible compared to the coalescent time scale, thus resulting merely in a (time-change) of a Kingman coalescent - a ‘weak’ seedbank effect.

## A Wright-Fisher nodel with geonetric seedbank distribution

We now introduce a Wright-Fisher type population model with mutation and seedbank in which individuals stay dormant for geometrically distributed amounts of time. The model is very much in line with classical probabilistic population genetics thinking (in particular assuming constant population size), but also captures several features of microbial seedbanks described in LENNON and JONES (2011), in particular reversible states of dormancy and mortality in the seedbank. We assume that the following (idealised) aspects of (microbial) dormancy can be observed:

(i) Dormancy generates a seedbank consisting of a reservoir of dormant individuals.
(ii) The size of the seedbank is comparable to the order of the total population size, say in a constant ratio *K*: 1 for some *K >* 0.
(iii) The size of the active population *N* and of the seedbank *M* = *M* (*N*) stays constant in time; combined with (ii) we get *N* = *K > M*.
(iv) The model is selectively neutral so that reproduction is entirely symmetric for all individuals; for concreteness we assume reproduction according to the Wright-Fisher mechanism in fixed generations. That means, the joint offspring distribution of the parents in each generation is symmetric multinomial. We interpret 0 offspring as the death of the parent, one offspring as mere survival of the parent, and two or more offspring as successful reproduction leading to new individuals created by the parent.
(v) Mutations may happen in the active population, at constant probability of the order *θ*_1_*/*(2*N*), but potentially also in the dormant population (at the same, or a reduced, or vanishing, probability *θ*_2_*/*(2*N*))
(vi) There is bi-directional and potentially repeated switching from active to dormant states, which appears essentially independently among individuals (‘spontaneous switching’). The individual initiation probability of dormancy per generation is of the order *c/N*, for *c >* 0.
(vii) Dormant individuals may die in the seedbank (due to maintenance and energy costs). If mortality is assumed to be positive, the individual probability of death per generation is of order *d/N*.
(viii) For each new generation, all these mechanisms occur independently of the previous generations.

We schematically visualise this mechanism in Figure 2, which is similar to Figure 1 in JONES and LENNON (2010). Vhether these assumptions are met of course needs to be determined for the concrete underlying real population. In this theoretical paper, we use the above assumptions to construct an explicit mathematical model that leads, measuring time in units of *N*, to a seedbank coalescent with mutation. Still, we wish to emphasise that, as dicussed in the previous section, the seedbank coalescent is robust as long as certain basic assumptions are met.

We now turn the above features into a formal mathematical model that can be rigorously analysed, extending the Wright-Fisher model with geometric seedbank component in BLATH *et al.* (2015b) by additionally including mortality in the seedbank and potentially different mutation rates in the active and dormant populations.

### Definition 1

(Seed bank model with mutation and mortality). Let *N ∊* ℕ, and let *c, K, θ*_1_ *>* 0 and *θ*_2_*, d ≥* 0. The seedbank model with mutation is obtained by iterating the following dynamics for each discrete generation *k ∊* ℕ_0_ (with the convention that all occuring numbers are integers; if not one may enforce this using appropriate Gauss brackets):

- The *N* active individuals from generation *k* = 0 produce 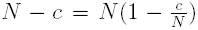 active individuals in generation *k* = 1 by multinomial sampling with equal weights.
- Additionally, *c* dormant individuals, sampled uniformly at random without replacement from the seedbank of size *M*:= *N/K* in generation 0, reactivate, that is, they turn into exactly one active individual in generation *k* = 1 each, and leave the seedbank.
- The active individuals from generation 0 are thus replaced by these (*N - c*) + *c* = *N* new active individuals, forming the active population in the next generation *k* = 1.
- In the seedbank, *d* individuals, sampled uniformly at random without replacement from generation *k* = 0, die.
- To replace the *c*+*d* vacancies in the seedbank, the *N* active individuals from generation 0 produce *c*+*d* seeds by multinomial sampling with equal weights, filling the vacant slots of the seeds that were activated.
- The remaining 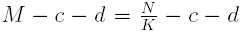. seeds from generation 0 remain inactive and stay in the seedbank
- During reproduction, each newly created individual copies its genetic type from its parent.
- In each generation, each active individual is affected by a mutation with probability *θ*_1_*/N,* and each dormant individual mutates with probability *θ*_2_*/N* (where *θ*_2_ may be 0).

This model is an extension of the model in BLATH *et al.* (2015b) to additionally include mortality in the seedbank and incorporate (potentially distinct) mutation rates in the active and dormant population. It appears to be a rather natural extension of the classical Wright-Fisher model. Note that the model has a geometric seedbank age distribution, since every dormant individual in each generation has the same probability to become active resp. die in the next generation, so that the time that an individual is in the dormant state is geometrically distributed. The parameter of this geometric distribution is given by

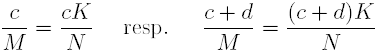

in the absence resp. presence of mortality in the seedbank. Vith mathematical arguments similar to those applied in BLATH *et al.* (2015b), it is now standard to show that the ancestral process of a sample taken from the above population model converges, on the coalescent time scale *N*, to the seedbank coalescent with parameters *c* and *K,* resp.

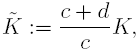

and mutation rates *θ*_1_*, θ*_2_. It is interesting to see that mortality leads to a decrease of the relative seedbank size in a way that depends on the initiation rate *c*, which is of course rather intuitive. In this sense 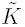 gives the ‘effective’ relative seedbank size.

### The type-frequencies in the bi-allelic seedbank population model

In this paper, we will mostly consider the *infinite sites model* (WATTEKSON, 1975), where it is assumed that each mutation generates an entirely new type. However, before turning to the infinite-site model, we briefly discuss the bi-allelic case, say with types {*a, A*}. Given initial type configurations *ξ*_0_ *∊* {*a, A*}^N^ and *η*_0_ *∊* {*a, A*}^M^, denote by

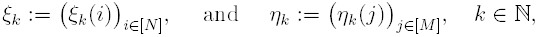

the genetic type configuration of the active individuals (*ξ*) and the dormant individuals (*η*) in generation *k* (obtained from the above mechanism). We assume that each mutation causes a transition from *a* to *A* or from *A* to *a.* Let

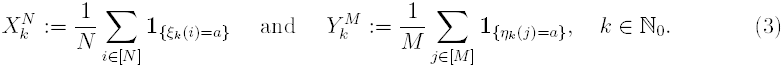

We call the discrete-time Markov chain 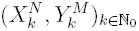 the *Wright-Fisher frequency process with mutation and seedbank component*. It can be seen from a generator computation that under our assumption with time measured in units of the active population size *N* it converges as *N → ∞* to the two-dimensional diffusion (*X*_*t*_, *Y*_*t*_)_*t*≥0_ that is the solution to the system of stochastic differential equations

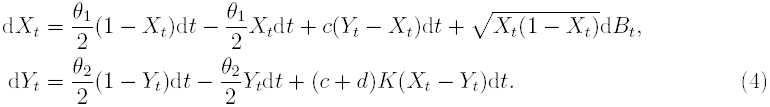

Here, (*B*_*t*_)_*t≥*0_ denotes standard one-dimensional Brownian motion. An alternative way to represent this stochastic process is via its Kolmogorov backward generator, cf. e. g. KARLIN and TAYLOR (1981), which is given by

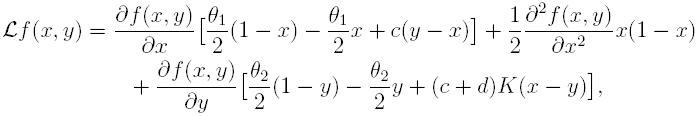

for functions *f ∊ C*^2^([0, 1]^2^). Existence and uniqueness of the stationary distribution of this process follows from compactness of the state space and strictly positive mutation rate *θ*__1__ *>* 0 in the active population. See Chapter 4.9 in ETHIER and KURTZ (2005) for more detailed arguments. Note that ℒ is reminiscent of the backward generator of the structured coalescent with two islands (HERBOTS, 1997; NOTOHARA, 1990); however, its qualitative behaviour is very different. Its relation to the structured coalescent with two islands will be investigated in future research. Observe that the solution of (4) is driven by only one Brownian motion.

## Population genetics with the seedbank coalescent

In contrast to LENNON and JONES (2011), who use a deterministic population dynamics approach to study seedbanks, we are interested in probabilistic effects of seedbanks on genetic variability. We therefore use a coalescent approach to study the (random) gene genealogy of a sample. In order to better understand how seedbanks shape genealogies, we consider genealogical properties, such as the time to most recent common ancestor, the total tree size, and the length of external branches.

### Genealogical tree properties

First we discuss some classical population genetic properties of the seedbank coalescent when viewed as a random tree without mutations. For the results that we derive below, it will usually be sufficient to consider the *block-counting process* (*N*_*t*_, *M*_*t*_)_*t≥*0_, of our coalescent, where *N*_*t*_ gives the number of lines in our coalescent that are active and *M*_*t*_ denotes the number of dormant lines *t* time units in the past. Then, (*N*_*t*_, *M*_*t*_)_*t≥*0_ is the continuous time Markov chain started in (*N*_0_, *M*_0_) ∊ ℕ_0_ × ℕ_0_ with transitions

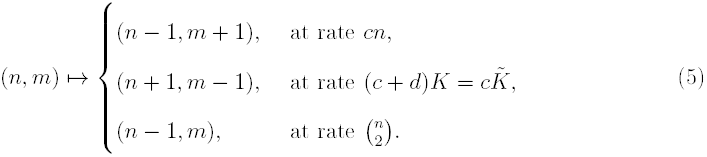

Again, introducing mutation can be done in the usual way, by superimposing independent Poisson processes with rate *θ*_1_/2 on the active lines, and at rate *θ*_2_/2 on the dormant lines. If the block-counting process is currently in state (*N*_*t*_, *M*_*t*_) = (*n, m*), then a mutation in an active line happens at rate *nθ*_1_/2, and a mutation in a dormant line at rate *mθ*_2_*/2.* The total jump rate from state (*n, m*) of the backward process with mutation is thus given by

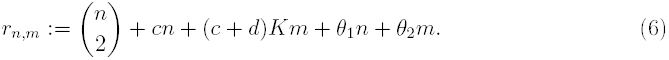

### Time to the most recent common ancestor

It has been shown in BLATH *et al.* (2015b) [Theorem 4.6] that the expected time to the most recent common ancestor (𝔼_*n*,0_[*T*_MRCA_]) for the seedbank coalescent, if started in a sample of active individuals of size *n*, is *O*(log log *n*), in stark contrast to the corresponding quantity for the classical Kingman coalescent, which is bounded by 2, uniformly in *n*, cf. (1). This already indicates that one should expect elevated levels of (old) genetic variability under the seedbank coalescent, since more (old) mutations can be accumulated. While the above result shows the asymptotic behaviour of the 𝔼_*n*,0_[*T*_MRCA_] for large *n*, it does not give precise information for the exact absolute value, in particular for ‘small to medium’ *n*. Here, we provide recursions for its expected value and variance that can be computed efficiently. First, we introduce some notation.

We define the *time to the most recent common ancestor* of the seedbank coalescent formally to be

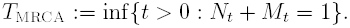

If the sample consists of *an* active and *bn* dormant individuals, for some *a, b ∊* ℝ^+^, then the expected time to the most recent common ancestor is log(*bn* + log *an*), (BLATH *et al.*, 2015b). Here, it is interesting to note that the time to the most recent common ancestor of the Bolhausen-Sznitman coalescent is also *O*(log log *n*) (GOLDSCHMIDT and MARTIN, 2005). The Bolthausen-Sznitman coalescent is often used as a model for selection, cf. e.g. NEHER and HALLATSCHEK (2013).

One can compute the expected time to most recent common ancestor recursively as follows. For *n, m ∊* ℕ_0_ let

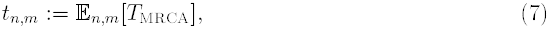

where 𝔼_*n*,*m*_ denotes expectation when started in (*N*_0_*, M*_0_) = (*n, m*), ie. with *n* active lines and *m* dormant ones. Observe that we need to consider both types of lines in order to calculate *t*_*n*,*m*_. Write

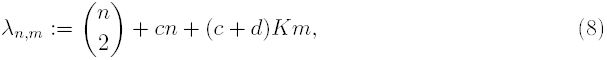

and abbreviate

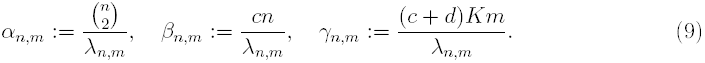

Then we have the following recursive representation

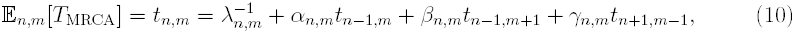

with initial conditions *t*_1,0_ = *t*_0,1_ = 0. The proof of (10) and a recursion for the variance of *T*_MRCA_ is given in Section S1. Since the process *N*_*t*_ + *M*_*t*_ is non-increasing in *t*, these recursions can be solved iteratively. In fact,

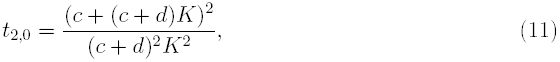

which in the case without mortality (*d* = 0) reduces to

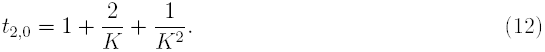

Notably, *t*__2,0__ is constant for sample size 2 (see Eq. 11) as *c* varies (Table 1) if *d* = 0, and in particular does not converge for *c →* 0 to the Kingman case. This effect is similar to the corresponding behaviour of the structured coalescent with two islands if the migration rate goes to 0, cf. NATH and GRIFFITHS (1993). However, the Kingman coalescent values are recovered as the seedbank size decreases (e.g. for *K* = 100 in Table 1).

**Table 1:** The expected time to most recent common ancestor (𝔼_*n*,0_ [*T*_MRCA_]) of the seedbank coalescent, obtained from (10), with seedbank size *K*, sample size *n*, dormancy initiation rates *c* as shown, and *d* = 0. All sampled lines are from the active population (sample configuration (*n,* 0)). For comparison, 𝔼_(*n*)_ [*T*_MRCA_] = 2(1 – 1*/n*) when associated with the Kingman coalescent (*K* = ∞). The multiplication × 10^4^ only applies to the first table with *K* = 0.01.

| $K = 0.01, \times 10^4$ | | | |
| --- | --- | --- | --- |
| $c$ | sample size $n$ | | |
|  | 2 | 10 | 100 |
| 0.01 | 1.02 | 2.868 | 5.185 |
| 0.1 | 1.02 | 2.731 | 4.487 |
| 1 | 1.02 | 2.187 | 2.666 |
| 10 | 1.02 | 1.878 | 2.085 |
| 100 | 1.02 | 1.84 | 2.026 |
| $K = 1$ | | | |
| $c$ | sample size $n$ | | |
|  | 2 | 10 | 100 |
| 0.01 | 4 | 10.21 | 17.18 |
| 0.1 | 4 | 9.671 | 14.97 |
| 1 | 4 | 8.071 | 10.02 |
| 10 | 4 | 7.317 | 8.221 |
| 100 | 4 | 7.212 | 7.954 |
| $K = 100$ | | | |
| $c$ | sample size $n$ | | |
|  | 2 | 10 | 100 |
| 0.01 | 1.02 | 1.846 | 2.052 |
| 0.1 | 1.02 | 1.838 | 2.026 |
| 1 | 1.02 | 1.836 | 2.02 |
| 10 | 1.02 | 1.836 | 2.02 |
| 100 | 1.02 | 1.836 | 2.02 |
| $K = \infty$ | 1 | 1.80 | 1.98 |

The fact that *t*__2,0__ = 4 for *K* = 1*, d* = 0 can be understood heuristically if *c* is large: In that situation, transitions between active and dormant states happen very fast, thus at any given time the probability that a line is active is about 1/2, and therefore the probability that both lines of a given pair are active (and thus able to merge) is approximately 1/4. We can therefore conjecture that for *d* = 0*, K* = 1 and *c → ∞* the genealogy of a sample is given by a time change by a factor 4 of Kingman’s coalescent.

Tables 1 and 2 show values of *t*_*n*,0_ obtained from (10) for various parameter choices and sample sizes. The relative size of the seedbank (*K*) has a significant effect on 𝔼_*n*,0_ [*T*_MRCA_]; a large seedbank (*K* small) increases 𝔼_*n*,0_ [*T*_MRCA_], while the effect of *c* is to dampen the increase in 𝔼_*n*,0_ [*T*_MRCA_] with sample size (Table (1)). The effect of the seedbank death rate *d* on 𝔼_*n*,0_ [*T*_MRCA_] is to dampen the effect of the relative size (*K*) of the seedbank (Table 2).

**Table 2:**
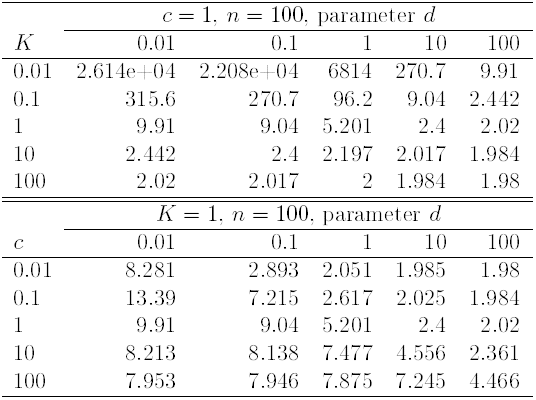
The expected time to most recent common ancestor (𝔼_*n*,0_ [*T*_MRCA_]) of the seed-bank coalescent, obtained from (10), with all *n* = 100 sampled lines assumed active, *c*, *K*, and *d* as shown. For comparison, 𝔼_(*n*)_ [*T*_MRCA_] = 2(1 – 1*/n*) (1.98 for *n* = 100) when associated with the Kingman coalescent.

### Total tree length and length of external branches

In order to investigate the genetic variability of a sample, in terms e.g. of the number of segregating sites and the number of singletons, it is useful to have information about the total tree length and the total length of external branches. Let *L*^(a)^ denote the total length of all branches while they are active, and *L*^(d)^ the total length of all branches while they are dormant. Their expectations

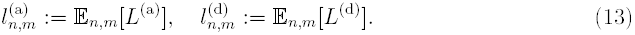

may be calculated using the following recursions for *n, m* ∊ ℕ__0__, and with *λ*_*n*,*m*_ given by (8),

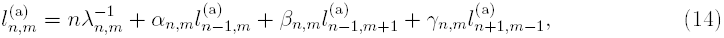

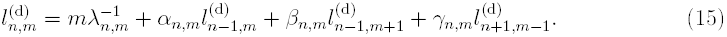

Observe that equations (14) and (15) differ in the factor (*n* resp. *m*) which multiplies 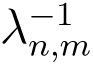. Similar recursions hold for their variances as well as for the corresponding values of the total length of external branches, which can be found in the Supplementary Information together with the respective proofs. From (14) and (15) one readily obtains

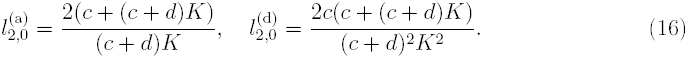

We observe that 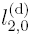 and 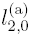 given in (16) are independent of *c* if *d* = 0 as also seen for *t*_2,0_ cf. (11). We will use (16) to obtain closed-form expressions for expected average number of pairwise differences.

The numerical solutions of (14) and (15) indicate that for *n* ≥2,

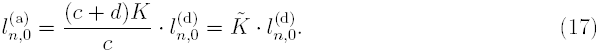

Hence, the expected total length of the active and the dormant parts of the tree are proportional, and ratio is given by the effective relative seedbank size.

Recursions for the expected total length of external branches are given in Prop. S1.3 in Supporting Information. Let 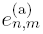 and 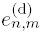 denote the expected total lengths of *active* and *dormant* external branches, respectively, when started with *n* active and *m* dormant lines. The numerical solutions of the recursions indicate that the ratio of expected values 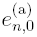 and 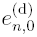 is also given by (17).

Recursions for expected branch lengths associated with any other class than singletons are more complicated to derive, and we postpone those for further study. Simulation results (not shown) suggest that the result (17) we obtained for relative expected total length of active branches, and active external branches, holds for all branch length classes; if 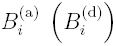 denotes the total length of *active (dormant)* branches subtending *i* ∊ {1, 2, …, *n* − 1} leaves, then, if all our sampled lines are active, we claim that 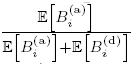 is given by (17).

Table S1 shows values of 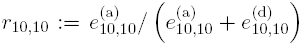, ie. the relative expected total length of external branches when our sample consists of ten active lines, and ten dormant ones. In contrast to the case when all sampled lines are active, *c* clearly impacts *r*__10,10__ when *d* is small. In line with previous results, *d* reduces the effect of the relative size (*K*) of the seedbank.

Table S2 shows the expected total lengths of active and dormant external branches 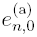 and 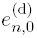 and for values of *c*, *K*, and *d* as shown. When the seedbank is large (*K* small), 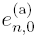 and 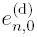 can be much longer than the expected length equal to 2 when associated with the Kingman coalescent (FU, 1995). However, as noted before, the effect of *K* depends on *d*. The effect of *c* also depends on *d*; changes in *c* have bigger effect when *d* is large.

One can gain insight into the effects of a seedbank on the site frequency spectrum by studying the effects of a seedbank on relative branch lengths. Let 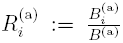 denote the relative total length of *active* branches subtending *i* leaves 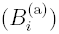, relative to the total length of active branches 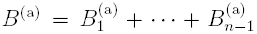, and we only consider the case when all *n* sampled lines are active. Thus, if one assumes that the mutation rate in the seedbank is negligible compared to the mutation rate in the active population, 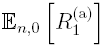 should be a good indicator of the relative number of singletons, relative to the total number of segregating sites. In addition, we investigate 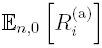 to learn if and how the presence of a seedbank affects genetic variation, even if no mutations occur in the seedbank. Figure S1 shows estimates of 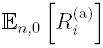 (obtained by simulations) for values of *c*, *K*, and *d* as shown (all *n* = 100 sampled lines assumed active). The main conclusion is that a large seedbank reduces the relative length of external branches, and increases the relative magnitude of the right tail of the branch length spectrum. Thus, one would expect to see a similar pattern in neutral genetic variation: a reduced relative number of singletons (relative to the total number of polymorphic sites), and a relative increase in the number of polymorphic sites in high count.

### Neutral genetic variation

In this subsection we derive and study several recursions for common measures of DNA sequence variation in the infinite sites model (ISM) of WATTERSON (1975). Samples are assumed to be drawn from the stationary distribution. We will also investigate how these quantities differ from the corresponding values under the Kingman coalescent, in an effort to understand how seedbank parameters affect genetic variability.

#### Segregating sites

First we consider the *number of segregating sites S in a sample*, which, assuming the ISM, is the total number of mutations that occur in the genealogy of the sample until the time of its most recent common ancestor. In addition to being of interest on its own, *S* is a key ingredient in commonly employed distance statistics such as those of TAJIMA (1989) and Fu and Li (1993). We let mutations occur on active branch lengths according to independent Poisson processes each with rate *θ*_1_/2, and on dormant branches with rate *θ*_2_/2. The expected value of *S* can be expressed in terms of the expected total tree-lengths as

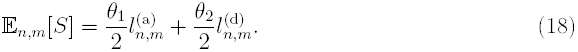

An alternative recursion for the expectation and the variance of the number of segregating sites, can be found in the supplementary material (Prop. S1.5).

Table 3 shows the expected number of segregating sites 𝔼_*n*,0_[*S*] = *s*_*n*,0_ in a sample of size *n* taken from the active population for values of *c* and *K* as shown. The size of the seedbank *K* strongly influences the number of segregating sites. If there is no mutation in the seedbank, it roughly doubles for *K* = 1 and approaches the normal value of the Kingman coalescent for small seedbanks (*K* = 100). The parameter *K* seems to have a more significant influence than the parameter *c*.

**Table 3:** The expected total number of segregating sites (*s*_*n*,0_), with values of *K*, *c*, *d* as shown, and sample size *n* = 100 (all lines from the active population); with *θ*_1_ = 2, and *θ*_2_ = 0. When associated with the Kingman coalescent, with *θ* = 2, and *s*_(100)_ = 10.35.

| values of $K$, $d = \theta_2 = 0$ | | | | | |
| --- | --- | --- | --- | --- | --- |
| $c$ | 0.01 | 0.1 | 1 | 10 | 100 |
| 0.01 | 1035 | 112.8 | 20.6 | 11.38 | 10.46 |
| 0.1 | 958.3 | 105.3 | 19.98 | 11.36 | 10.46 |
| 1 | 790.6 | 90.37 | 19.4 | 11.38 | 10.46 |
| 10 | 884 | 99.92 | 20.16 | 11.39 | 10.46 |
| 100 | 1010 | 110.8 | 20.61 | 11.39 | 10.46 |
| values of $K$, $d = 100$, $\theta_2 = 0$ | | | | | |
| $c$ | 0.01 | 0.1 | 1 | 10 | 100 |
| 0.01 | 10.46 | 10.37 | 10.36 | 10.35 | 10.35 |
| 0.1 | 11.36 | 10.46 | 10.37 | 10.36 | 10.35 |
| 1 | 19.32 | 11.37 | 10.46 | 10.37 | 10.36 |
| 10 | 91.97 | 19.3 | 11.29 | 10.45 | 10.36 |
| 100 | 510.3 | 60.83 | 15.51 | 10.87 | 10.41 |

#### Average pairwise differences

Average pairwise differences are a key ingredient in the distance statistics of TAJIMA (1983) and FAY and WU (2000). Expected value and variance for average pairwise differences in the Kingman coalescent were first derived in TAJIMA (1983). Here, we give an expression for the expectation in terms of the expected total tree lengths. Denote by *π* the average number of pairwise differences

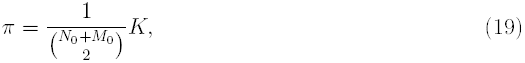

where 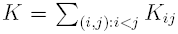 is the total number of pairwise differences, with *K*_*ij*_ denoting the number of differences observed in the pair of DNA sequences indexed by (*i, j*). We abbreviate *d*_*n*,*m*_:= 𝔼_*n*,*m*_[*K*] and obtain

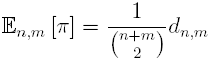

which can be calculated using

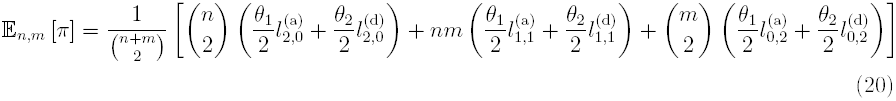

Where 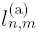 and 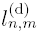 are defined in (13).

Hence, given a sample configuration (*n,* 0), i.e. our *n* sampled lines are all active, (20), together with (16), gives

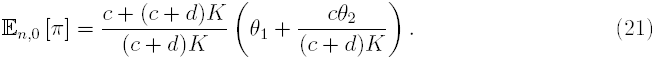

If now *d* = 0, the dependence on *c* disappears again, since we have

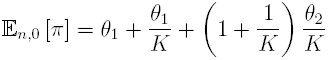

which is obviously highly elevated compared to *θ*_1_ if the seedbank is large (*K* small). For comparison, 𝔼_(*n*)_ [*π*] = *θ*_1_ when associated with the usual Kingman coalescent, which we recover in the absence of a seedbank (*K → ∞*) in (21).

#### The site-frequency spectrum (SFS)

The site frequency spectrum (SFS) is one of the most important summary statistics of population genetic data in the infinite sites model. Suppose that we can distinguish between mutant and wild-type, e.g. with the help of an outgroup. As before, we distinguish between the number of samples taken from the active population (say *n*) and the dormant population (say *m*). Then, the SFS of an (*n, m*)-sample is given by

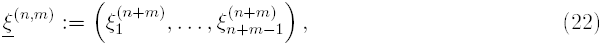

where the 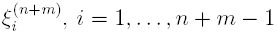 denote the number of sites at which variants appear *i*-times in our sample of size *n* + *m*. For the Kingman coalescent, the expected values, variances and covariances of the SFS have been derived by Fu (1995). Expected values and covariances can be computed in principle extending the theory in Fu (1995) resp. GRIFFITHS and TAVARÉ (1998), However, the computations would be far more involved than the previous recursions and will be treated in future research. We derive recursions for the expected number of singletons, and investigate the whole SFS by simulation.

#### Number of singletons

The number of singletons in a sample is often taken as an indicator of the kind of historical processes that have acted on the population. By ‘singletons’ we mean the number of derived (or new) mutations which appear only once in the sample, which in the infinite sites model, are equal to the number of mutations occurring on external branches. Thus we can relate the expected number of singletons, denoted by 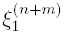, to the total length of external branches in the same way as we related the number of segregating sites to the total tree length. Let 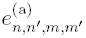 denote the expected total length of external branches when our sample consists of *n active external* lines, *n′ active internal* lines, *m dormant external* lines, and *m′ dormant internal* lines. Define 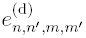 similarly as the expected total length of dormant external branches. Recursions for 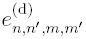 and 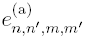 are given in the supplementary material. For *n, m ∊* ℕ_0_ we have that the expected number of singletons is given by

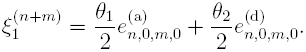

Thus, one can compute the expected number of singtetons by solving the recursions for external branch lengths. By way of example, Table S2 gives values of 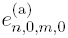 and 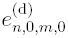 for a sample of 10 active lines (*n* = 10, *m* = 0).

#### The whole site-frequency spectrum

Figure 3 shows estimates of the normalised expected frequency spectrum 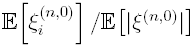, where *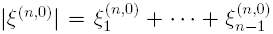* denotes the total number of segregating sites. Figure 3 shows that if the relative size of the seedbank is small (say, *K* = 100), then the SFS is almost unaffected by dormancy, in line with intuition. If the seedbank is large (say *K* = 0.1) and the transition rate *c* = 1 is comparable to the mutation rate *θ*_1_*/*2 = 1 then the spectrum differs significantly, in particular the number of singletons is reduced by about one-half, which should be significant, and the right-tail is much heavier.

**Figure 3:**
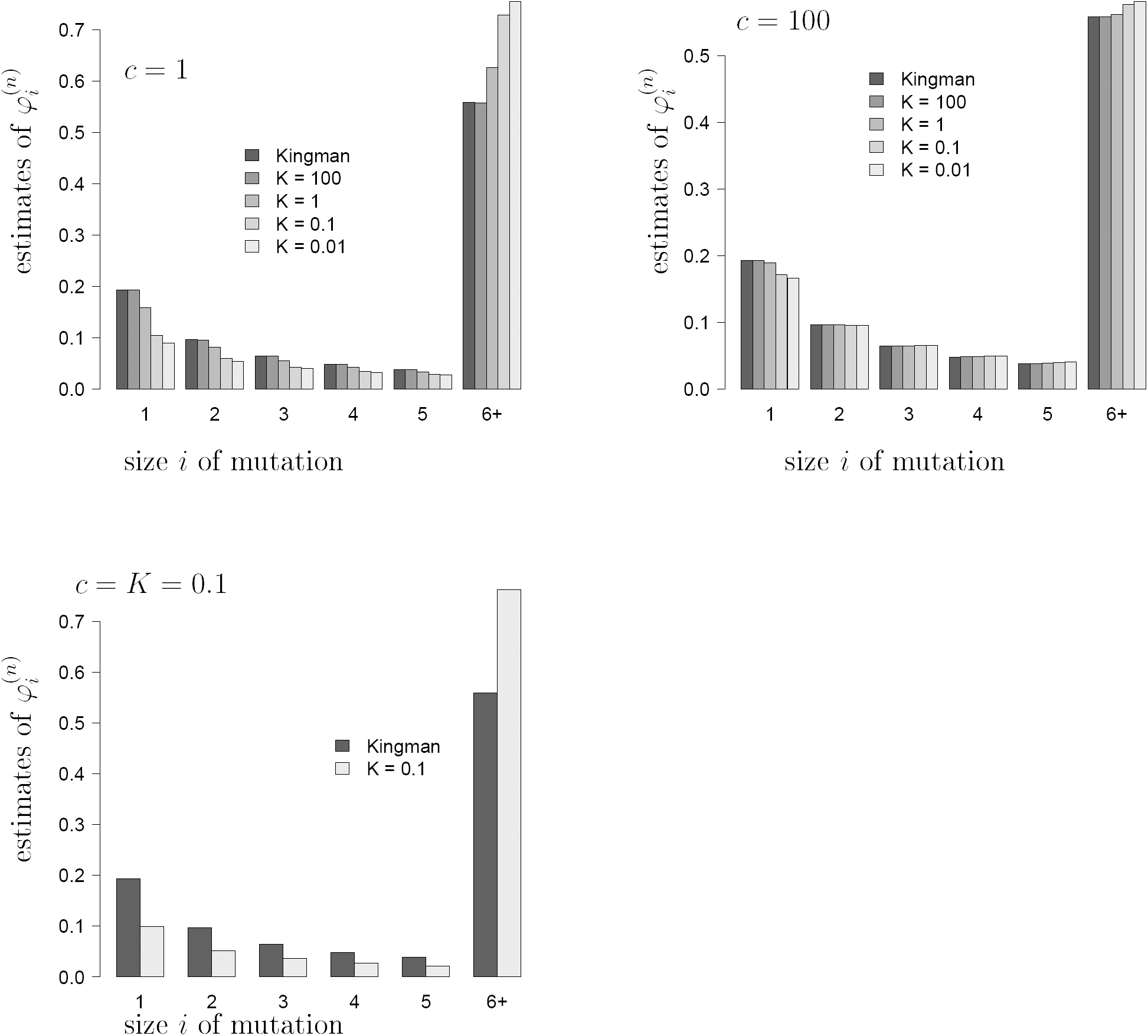
Estimates of the normalised site-frequency spectrum 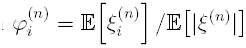 with all *n* = 100 sampled lines assumed active, and values of *c* and *K* as shown (*d* = 0). The mutation rate in the active population is fixed: *θ*_1_ = 2, and there is no mutation in the dormant states (*θ*_2_ = 0). All estimates based on 10^5^ replicates.

This can be understood as follows: if the seedbank leads to an extended time to the most recent common ancestor, then the proportion of old mutations should increase, and these should be visible in many sampled individuals, strengthening the right tail of the spectrum.

It is interesting to see that even in the presence of a large seedbank (say *K* = 0.1), large transitions rates (say *c* = 100) do not seem to affect the normalised spectrum. Again, this can be understood intuitively, since by the arguments presented in the discussion after (12) large *c* should lead to a constant time change of the Kingman coalescent (with a time change depending on *K*). Such a time change does not affect the normalised spectrum.

One reason for considering the SFS is naturally that one would like to be able to use the SFS in inference, to determine, say, if a seedbank is present, and how large it is. If one has expressions for the expected SFS under some coalescent model, one can use the normalised expected SFS in an approximate likelihood inference (see eg. ELDON *et al.*, 2015). The normalised spectrum is also appealing since it is quite robust to changes in the mutation rate (ELDON *et al.*, 2015). For comparison, Figure S2 shows estimates of the expected normalised spectrum 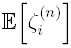 where *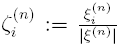,* and shows a similar pattern as for the normalised expected spectrum in Figure 3.

## Distance statistics

Rigorous inference work is beyond the scope of the current paper. However, we can still consider (by simulation) estimates of the distribution of various commonly employed distance statistics. Distance statistics for the site-frequency spectrum are often employed to make inference about historical processes acting on genetic variation in natural populations. Commonly used statistics include the ones of TAJIMA (1989) (*D*_T_), Fu and Li (1993) (*D*_FL_), and FAY and WU (2000) (*D*_FW_). These statistics contrast different parts of the site-frequency spectrum (cf. eg. ZENG *et al.*, 2006).

### The ℓ_2_ distance

Arguably the most natural distance statistic to consider is the ℓ_2_-distance (or sum of squares) of the whole SFS (or some lumped version thereof) between the observed SFS and an expected SFS based on some coalescent model. The 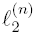 statistic (*n* denotes sample size) is given by

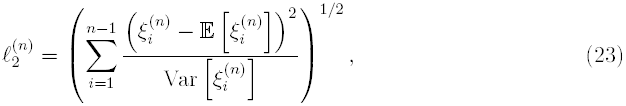

where, in our case, expectation and variance are taken with respect to the Kingman coalescent (Fu, 1995). Estimates of the distribution of 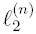 are shown in Figure 4. As the size of the seedbank increases (*K* decreases), one observes worse fit of the site-frequency spectrum with the expected SFS associated with the Kingman coalescent.

**Figure 4:**
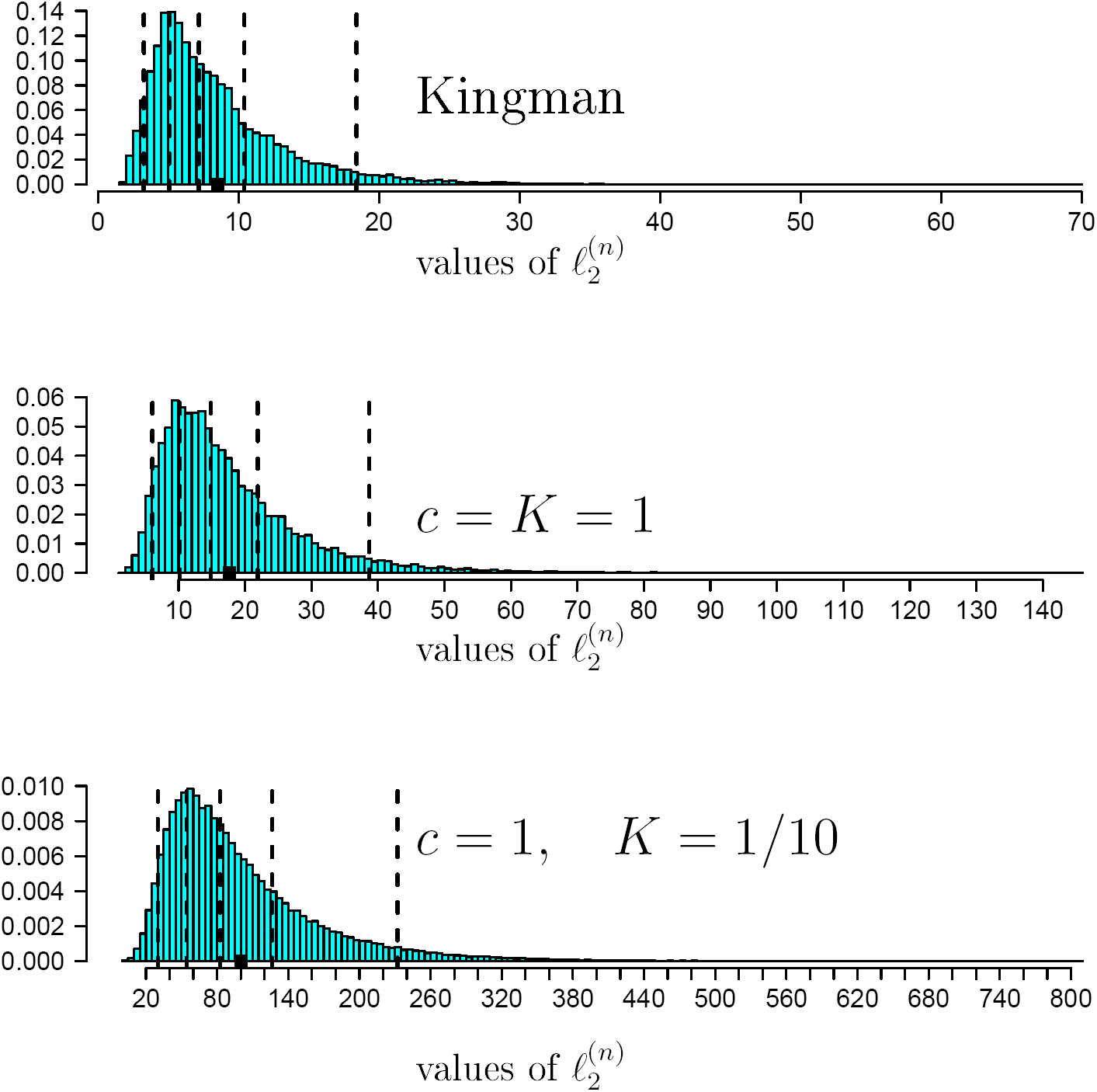
Estimates of the distribution of the 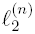 statistic (23), with all *n* = 100 sampled lines assumed active, *c* and *K* as shown, *θ*_1_ = 2, *θ*_2_ = 0. The vertical broken lines are the 5%, 25%, 50%, 75%, 95% quantiles and the black square (▀) denotes the mean. The entries are normalised to have unit mass 1. All estimates are based on 10^5^ replicates.

### Tajima's *D*

Tajima’s statistic (*D*_T_) for a sample of size *n*, with 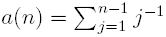, is defined as

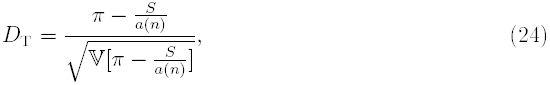

(TAJIMA, 1989) where the variance 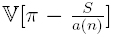 depends on the mutation rate *θ* which is usually estimated from the data. Under the Kingman coalescent, 𝔼[*D*_T_] = 0. Deviations from the Kingman coalescent model become significant at the 5% level if they are either greater than 2 or smaller than −2. Negative values of *D*_T_ should appear if there is an excess of either low- or high-frequency polymorphisms and deficiency of middle frequency polymophisms (see e.g. WAKELEY (2009) for further details). Positive values of *D*_T_ are to be expected if variation is common with moderate frequencies, for example in presence of a recent population bottleneck, or balancing selection.

The empirical distribution of *D*_T_ was investigated by simulation for different seedbank parameters (Figures 5, S3), assuming that mutations do not occur in the seedbank (*θ*_2_ = 0). If the seedbank is large (*K* = 1*/*10, 1*/*100), then the median of *D* becomes significantly positive. For *c* = *K* = 1, there is very little deviation from the Kingman coalescent. Again *D* seems to be more sensitive to small values of *K* than changes in *c*. This is in line with our results on the 𝔼_*n*,0_ [*T*_MRCA_], with highly elevated times for small *K*. In the latter case, old variation will dominate, thus resembling a population bottleneck, producing positive values of *D*_T_.

**Figure 5:**
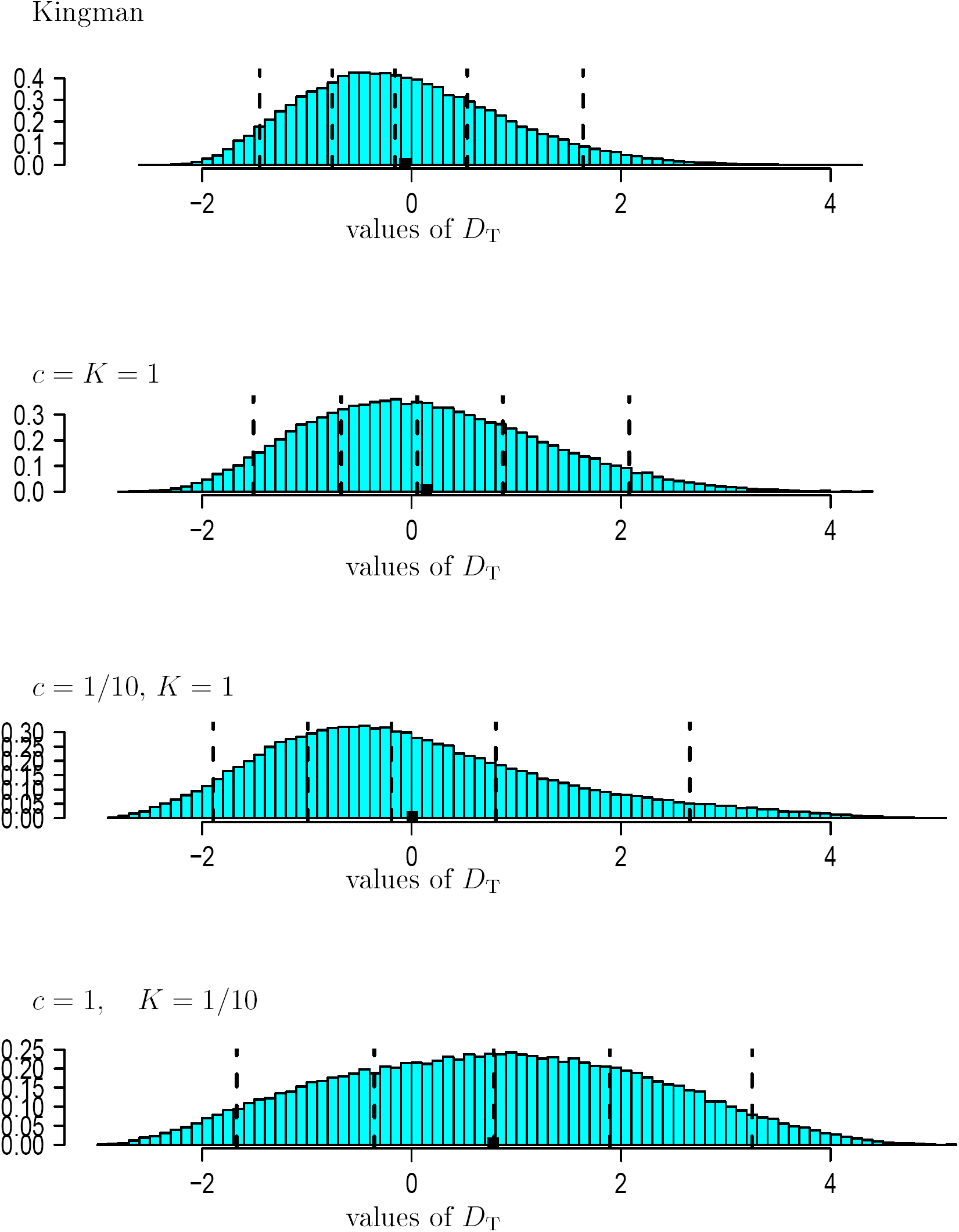
Estimates of the distribution of Tajima’s *D*_T_ (24) with all *n* = 100 sampled lines assumed active, *θ*_1_ = 2, *θ*_2_ = 0. The vertical broken lines are the 5%, 25%, 50%, 75%, 95% quantiles and the black square (▮) denotes the mean. The entries are normalised to have unit mass 1. The histograms are drawn on the same horizontal scale. Based on 10^5^ replicates.

In conclusion, *D*_T_ might not be a very good statistic to detect seedbanks.

### Fu and Li’s *D*

Fu and LI (1993) statistic *D*_FL_ is defined as

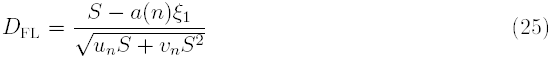

with *S* being the total number of segregating sites, *ξ*_1_ the total number of singletons, 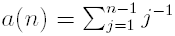, and *u*_*n*_ and *v*_*n*_ as in Fu and Li (1993) (see also Durrett 2008). As with *D*_T_, 𝔼[*D*_FL_] = 0 under the Kingman coalescent.

Figure 6 shows estimates of the distribution of *D*_FL_ assuming *θ*_2_ = 0. When the seedbank is large (*K* small), the distribution of *D*_FL_ becomes highly skewed, with most genealogies resulting in low number of singletons compared with the total number of polymorphisms, resulting in positive *D*_FL_. This is in line with our observations about the relative number of singletons associated with a large seedbank (Figures 3, S2), and the relative length of external branches (Figure S1).

**Figure 6:**
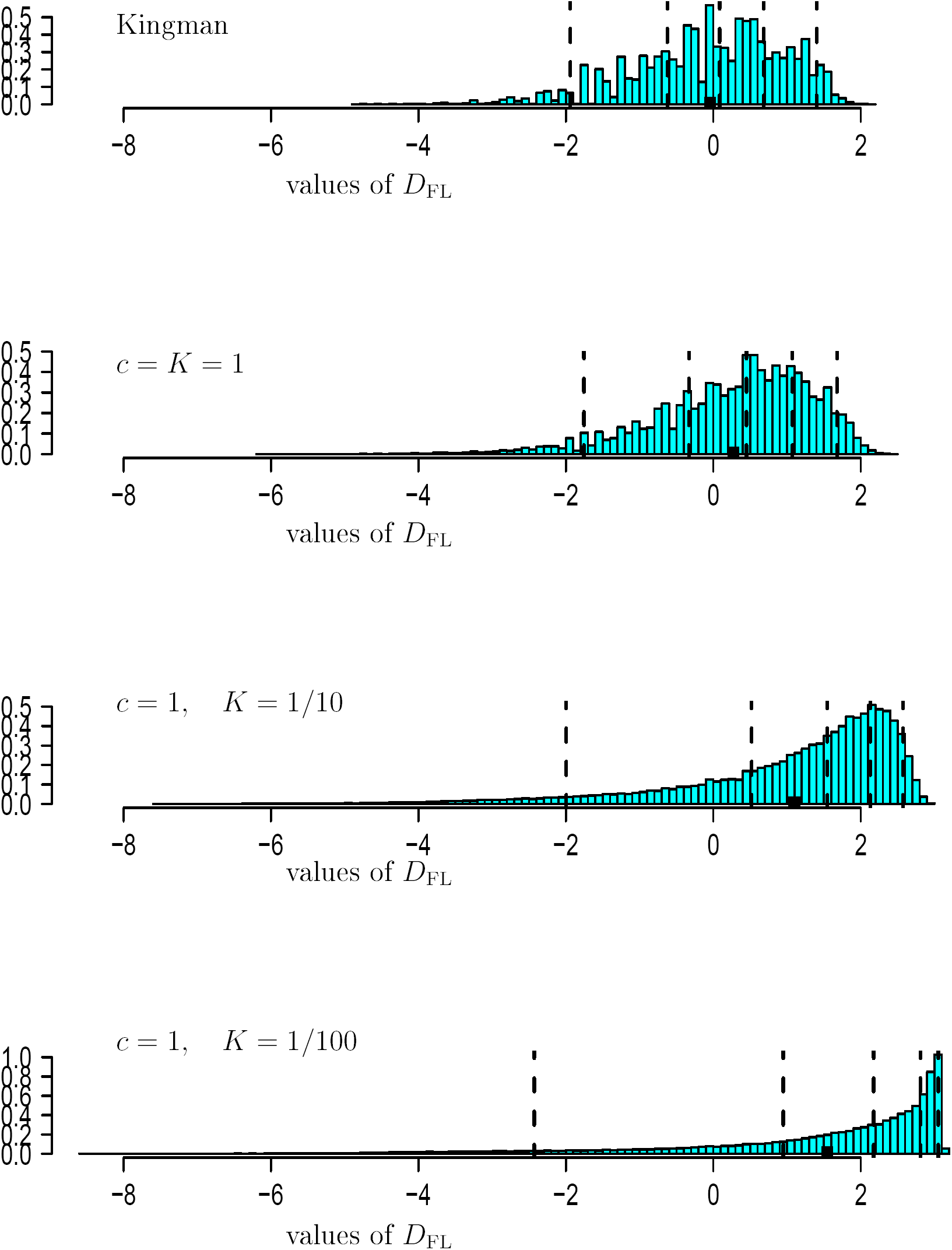
Estimates of the distribution of Fu and Li’s *D*_FL_ (25) with all *n* = 100 sampled lines assumed active, *θ*_1_ = 2, *θ*_2_ = 0. The vertical broken lines are the 5%, 25%, 50%, 75%, 95% quantiles and the black square (▮) denotes the mean. The entries are normalised to have unit mass 1. The histograms are drawn on the same horizontal scale. Based on 10^5^ replicates.

### Fay and Wu’s *H*

The distance statistics *D*_FW_ of FAY and WU (2000) is defined as

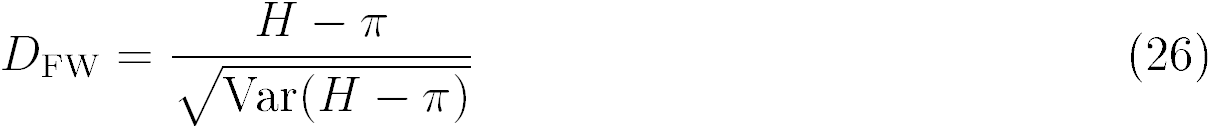

where

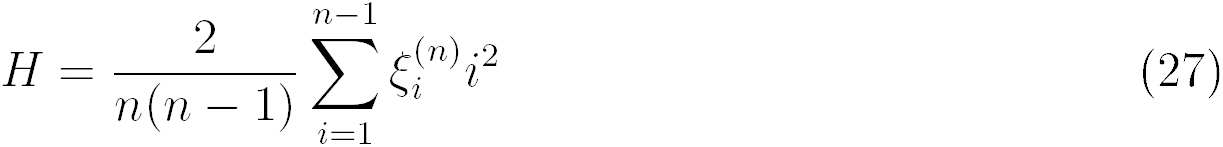

and *π* is the average number of pairwise differences. A formula for the variance of *D*_FW_was obtained by ZENG *et al.* (2006). Figure 7 holds estimates of the distribution of *D*_FW_with *n* = 100, *d* = 0, and *c* and *K* as shown. As the seedbank size increases (*K* decreases) high frequency variants, as captured by *H*, become dominant over the middle-frequency variants captured by *π*. In conclusion, Fu and Li’s *D*_FL_, or Fay and Wu’s *D*_FW_ may be preferrable over Tajima’s statistic *D*_T_ to detect the presence of a seedbank. A rigorous comparison of different statistics (including the *E* statistic of ZENG *et al.* (2006)), and their power to distinguish between absence and presence of a seedbank, must be the subject of future research.

The C code written for the computations is available at http://page.math.tu-berlin.de/∼eldon/programs.html.

**Figure 7:**
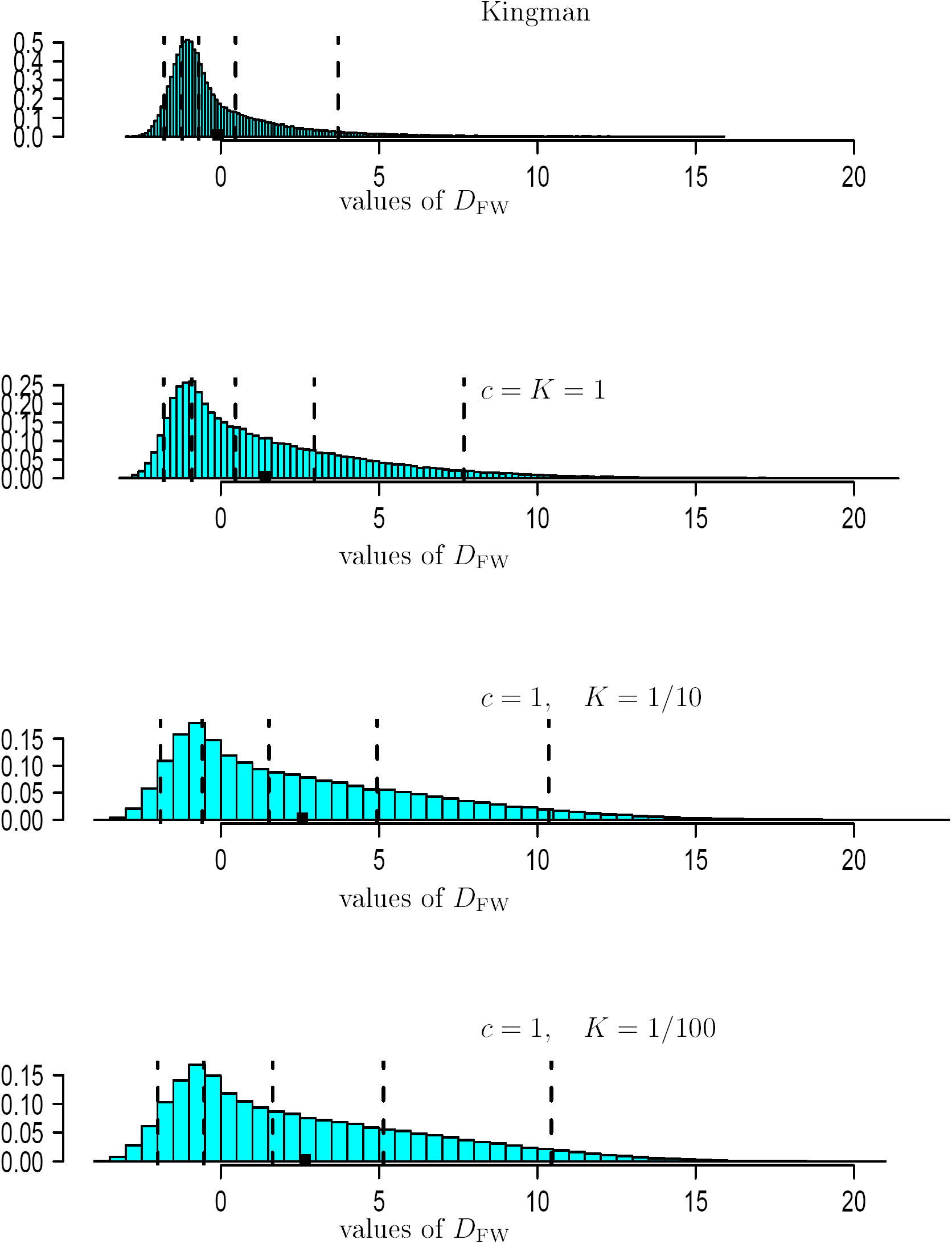
Estimates of the distribution of Fay and Wu’s *D*_FW_ (26) with all *n* = 100 sampled lines assumed active, *θ*_1_ = 2, *θ*_2_ = 0. The vertical broken lines are the 5%, 25%, 50%, 75%, 95% quantiles and the black square (▮) denotes the mean. The entries are normalised to have unit mass 1. The histograms are drawn on the same horizontal scale. Based on 10^5^ replicates.

## Discussion

In the previous sections, we have presented and analysed an idealised model of a population sustaining a large seedbank, as well as the resulting patterns of genetic variability, with the help of a new coalescent structure, called the seedbank coalescent (with mutation). This ancestral process appeared naturally as scaling limit of the genealogy of large populations producing dormant forms, in a similar way as the classical Kingman coalescent arises in conventional models, under the following assumptions: the seedbank size is of the same order as the size of the active population, the population and seedbank size is constant over time, and individuals enter the dormant state by spontaneous switching independently of each other, in a way that individual dormancy times are comparable to the active population size. We begin with a discussion of these modeling assumptions.

The assumption that the seedbank is of comparable size to the active population is based on LENNON and JONES (2011), where it is shown in Box 1, Table a, that this is often the case in microbial populations.

Assuming constant population size is a very common simplification in population genetics, and can be explained with constant environmental conditions. We claim that ‘weak ’ fluctuations (of smaller order than the active population size) still lead to the seedbank coalescent model, as is the case for the Kingman coalescent. However, seedbanks are often seen as a bet hedging strategy against drastic environmental changes, which is not yet covered by our models. We see this as an important task for future research, which will require serious mathematical analysis. In the case of weak seedbank effects, fluctuating population size has been considered in ŽIVKOVIĆ and TELLIER (2012), where the presence of the seedbank was observed to leading to an increase of the effective population size.

Assuming spontaneous switching of single individuals between active and dormant state is also based on LENNON and JONES (2011) [p. 122/124]. This is somewhat restricting the scope of the model because it will not capture major environmental changes that may trigger a simultaneous change of state of a large proportion of individuals (e.g. due to sudden lack of nutrients). This effect is closely related to drastic changes in population size, and again may lead to serious alterations of our predictions. Hence, including such large switching events will also be an important part of our future work (and will again require substantial mathematical work). In VITALIS *et al.* (2004) a whole proportion of the dormant population becomes active in every generation, but this should be seen in conjunction with the fact that dormancy is of limited duration, which excludes drastic alterations on a long time scale. Assuming that the time spent in the seedbank is of the order of the population size is one of the main features that distinguishes our model from previous models of weak seedbank effects as previously investigated in KAJ *et al.* (2001); VITALIS *et al.* (2004). Statistical inference will be needed to support or reject this assumption, and to distinguish between weak and strong seedbanks. One distinguishing feature of weak and strong seedbanks is the behaviour of the normalised site frequency spectrum. Since weak seedbanks lead to a genealogy which is a constant time change of Kingman’s coalescent KAJ *et al.* (2001); BLATH *et al.* (2013) the normalised frequency spectrum of weak seedbanks will be similar to those corresponding to the Kingman coalescent, while under our model we observe (at least for large seedbanks) a reduction in the number of singletons (Figure S2). The model of Kaj *et al.* (2001) was used in TELLIER *et al.* (2011), where Tajima’s D was used in order to detect seedbanks.

We now discuss our results for the behaviour of classical quantities describing genetic variability under our modeling assumptions, that is, when the genealogy of a sample can be described by the seedbank coalescent. In particular, we used it to derive recursions for quantities such as the time to the most recent common ancestor, the total tree length or the length of external branches. We investigated statistics of interest to genetic variability such as the number of segregating sites, the site frequency spectrum, Tajima’s *D*, Fu and Li’s *D* and Fay and Wu’s *H* by numerical solution of our recursions and by simulation. It turns out that the seedbank size *K* leads to significant changes for example in the site frequency spectrum, producing a positive Tajima’s *D*, indicating the presence of old genetic variability, in line with intuition. Interestingly, the the influence of *c* seems to be less pronounced. For *K → ∞* we observe convergence towards the Kingman coalescent regime, while *c → ∞* seems to lead to a constant time change of Kingman’s coalescent.

We are confident that our results so far have the potential to open up many interesting research questions, both on the modeling and on the statistical inference side, as well as in data analysis. For example, it should be interesting to derive a test to distinguish between the presence of strong vs. weak (resp. negligible) seedbanks. Another important task in future research will be to infer parameters of the model. While the relative seedbank size *K* can in principle be directly observed by cell counting (LENNON and JONES, 2011), the parameter *c* seems to be difficult to observe, in particular because we have seen that many statistics we calculated are independent of or at least not very sensitive with respect to *c.* On the other hand, this shows that our results are fairly robust under alterations of *c,* such that estimations or tests may be applied to some extent without prior knowledge on *c.* The mortality rate *d* may for many practical purposes be included into the parameter *K* or 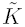 measuring the “effective” relative seedbank size.

Estimating the mutation rates *θ*_1_ and *θ*_2_ is another goal for the future. In particular, in view of an ongoing debate on the possibility of mutations in dormant individuals (MAUGHAN, 2007), it would be important to devise a test to determine if *θ*_2_ *>* 0.

## Acknowledgements

JB, BE, and NK acknowledge support by Deutsche Forschungsgemeinschaft (DFG) as part of SPP Priority Programme 1590’Probabilistic Structures in Evolution’; JB and BE through DFG grant BL 1105/3-1; JB and NK through grant DFG BL 1105/4-1. ACG is supported by DFG RTG 1845 ‘Stochastic Analysis and Applications in Biology, Finance, and Physics’, the Berlin Mathematical School (BMS), and the Mexican Council of Science (CONACyT) in collaboration with the German Academic Exchange Service (DAAD). MWB is supported by DFG RTG 1845, and the BMS.

## SUPPORTING INFORMATION

### S1 Proofs and further recursive formulas

#### Expectation and variance of the *T*MRCA

For *n, m ∊* ℕ_0_, let *t*_*n*,*m*_:= 𝔼_*n*,*m*_ [*T*_MRCA_] and *v*_*n*,*m*_:= V_*n*,*m*_[*T*_MRCA_].

##### Proposition S1.1.

Let *n, m ∊* ℕ_0_. *Then we have the following recursive representations*

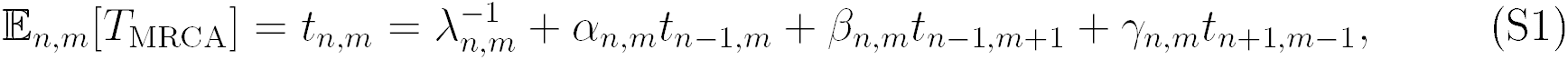

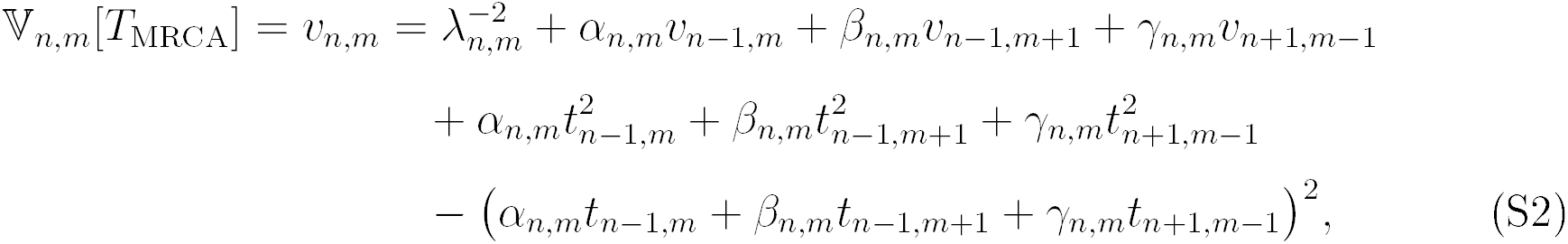

*with initial conditions t*_1,0_ = *t*_0,1_ = *v*_1,0_ = *v*_0,1_ = 0.

*Proof of Proposition S1.1.* Let *τ*_1_ denote the time of the first jump of the process (*N*_*t*_, *M*_*t*_)_*t≥*0_. If started at (*n, m*), this is an exponential random variable with parameter *λ*_*n*,*m*_. Applying the strong Markov property we obtain

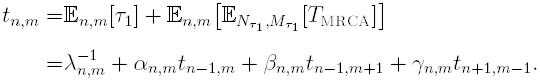

Similarly, the strong Markov property (telling us that *τ*_1_ is independent of the time to the most recent common ancestor of the (random) sample (*N*_*τ*1_, *M*_*τ*1_) and the law of total variance yields

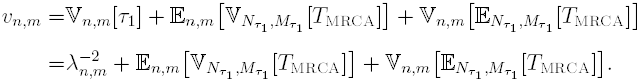

We have

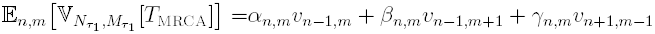

and

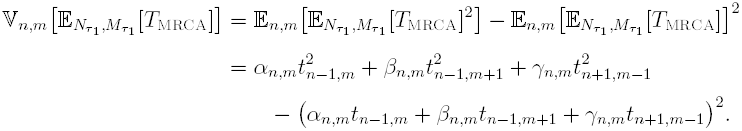

Combining the observations proves the result.

#### Expectation and variance of the total tree length

Let 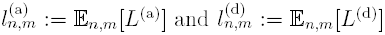 denote the expectations, and *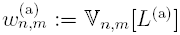* and 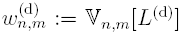 the variances of the total tree lengths, and define the mixed second moment, 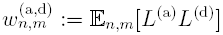.

#### Proposition S1.2

(Recursion: Total tree length). For *n, m ∊* ℕ *we have*

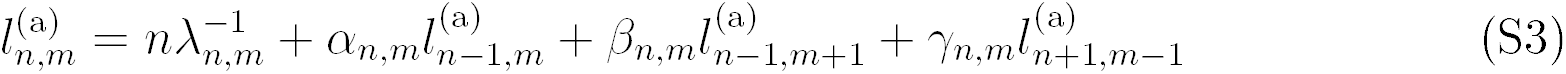

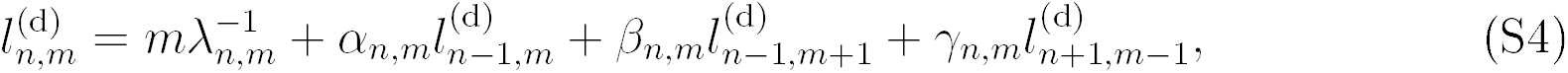

*and*

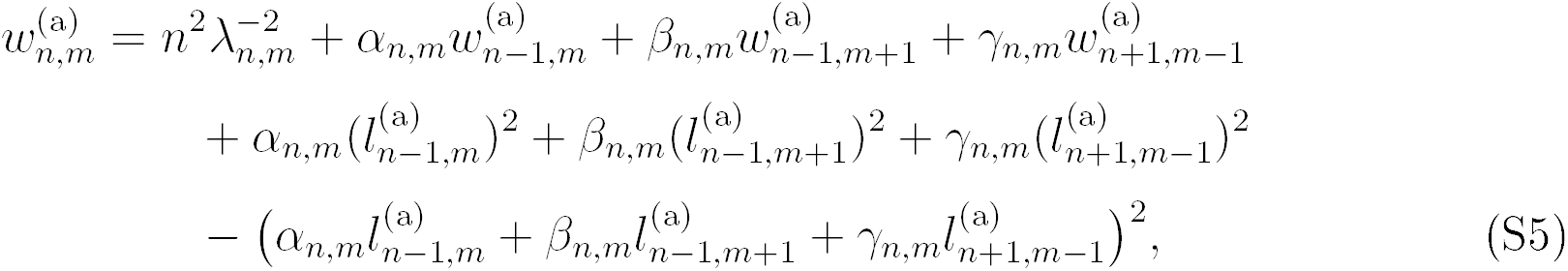

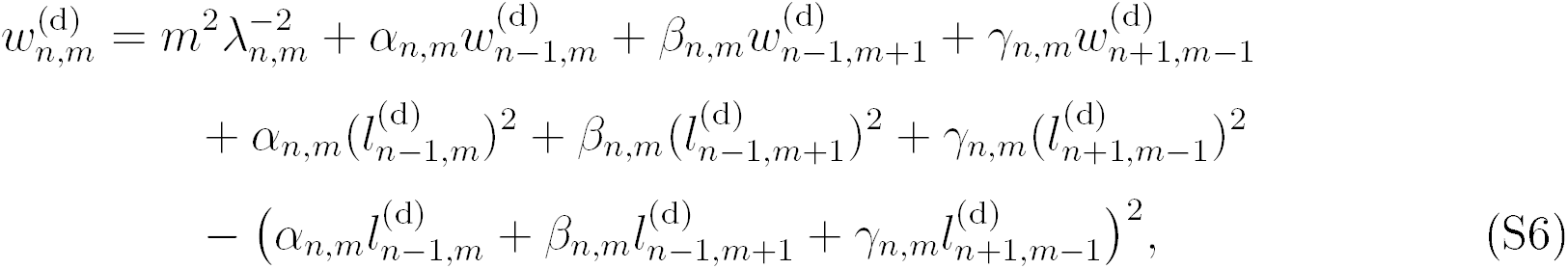

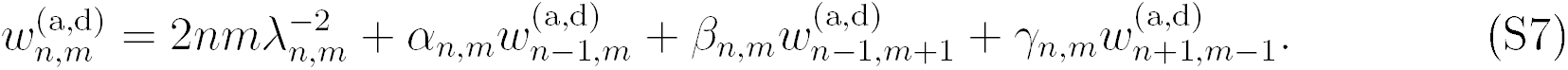

*Proof of Proposition S1.2.* The result can easily be obtained observing that each stretch of time of length *τ* in which we have a constant number of *n* active blocks and *m* dormant blocks contributes with *nτ* to the total active tree length, and with *mτ* to the total dormant tree length. Thus we have

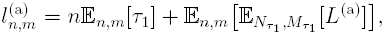

and we proceed as in the proof of Proposition S1.1. From these quantities we easily obtain the expected total tree length as *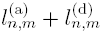.* Moreover,

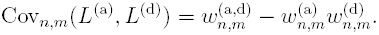

#### Expectation of total length of external branches

To derive recursions for the total length of external branches in either of the two states is a little more involved, since obviously a coalescence can happen between either two external active branches, two internal active branches, or an external and an internal active branch. We use indices (*n, n’, m, m’*) to denote the number of external active branches, internal active branches, external dormant branches, and internal dormant branches, respectively. Abbreviate

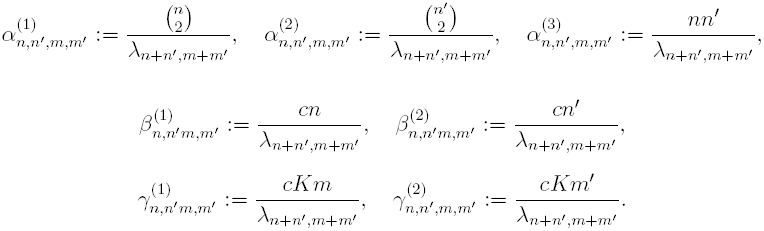

Let *E*^(a)^ denote the total length of external branches in the plant state, and *E*^(d)^ the total length of external branches in the seed state. Then we have

#### Proposition S1.3

(Recursion: Total length of external branches). For *n, m ∊* ℕ, *we have the representation*

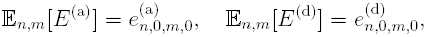

*where* 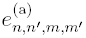 *and* 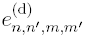n; n’; m;m’ ∊ ℕ_0_ *satisfy the recursions*

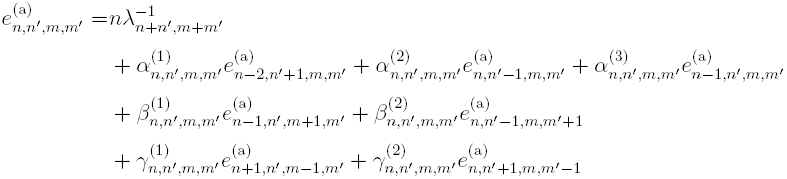

*and*

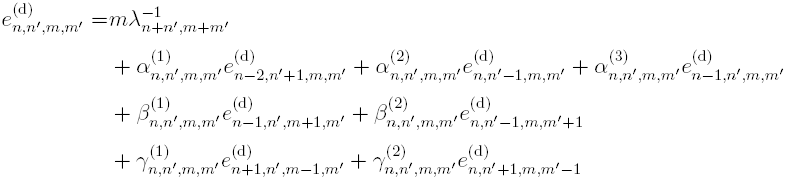

Observing that *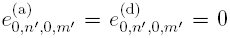* for all *n', m',* and 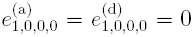 and that the total number *n* + *n'* + *m* + *m'* is non-increasing, these recursions can be solved iteratively.

*Proof of Proposition S1.3.* This follows by a similar first-step analysis as in Proposition S1.2, taking into account the transitions for internal and external branches, and observing that at each coalescence event between two external branches, the number of external plant branches is reduced by two and the number of internal branches is increased by one, in a coalescence of an external and an internal branch, the number of external plant branches is reduced by one and the number of internal plant branches stays the same, and in a coalescence of two internal branches, their number is reduced by one.

Obviously, the expected total length of external branches is then given by 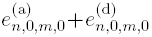

Note that proceeding as in Proposition S1.2, we could also give recursions for the variances of these quantities.

#### Expectation and variance of the number of segregating sites

##### Proposition S1.4

For *n, m ∊* ℕ_0_ we have

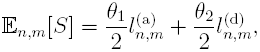

and

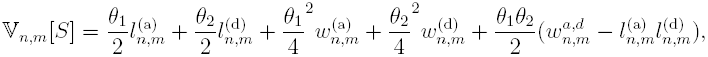

*where 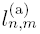, 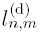,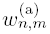, 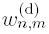 and 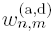 are given by Proposition S1.2.*

*Proof of Proposition S1.4.* Observe that conditional on the total lengths *L*^(a)^*, L*^(d)^, the number of segregating sites is the sum of two independent Poisson random variables with parameters *θ*_1_*L*^(a)^*/*2 and *θ*_2_*L*^(d)^*/*2, respectively. Hence, if an ancestral line is in the plant state for a period of time of length *L >* 0, the expected number of mutations that occur in this period is *L θ* _1_*/*2. Similarly, in a period of length *L* when the ancestral line is a seed, the expected number of mutations is *L θ* _2_*/*2. Thus the first result follows directly from Proposition S1.2.

For the second result, we apply the law of total variance and obtain similarly that

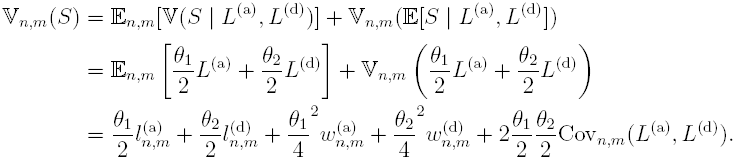

It is possible to directly derive a recursion for the number of segregating sites without explicitly passing through calculating the tree lengths. Since it may be of use we state it here. Let

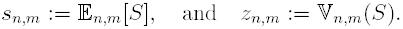

##### Proposition S1.5

(Alternative recursion). Let *n, m ∊* ℕ_0_. Then

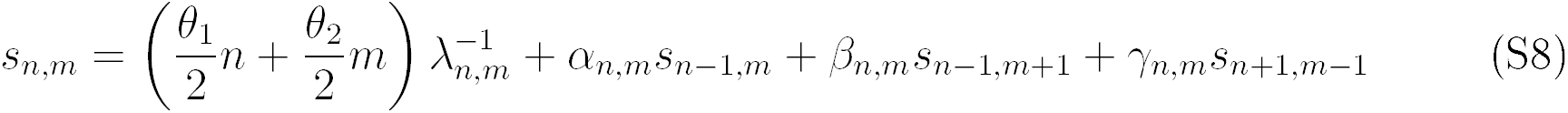

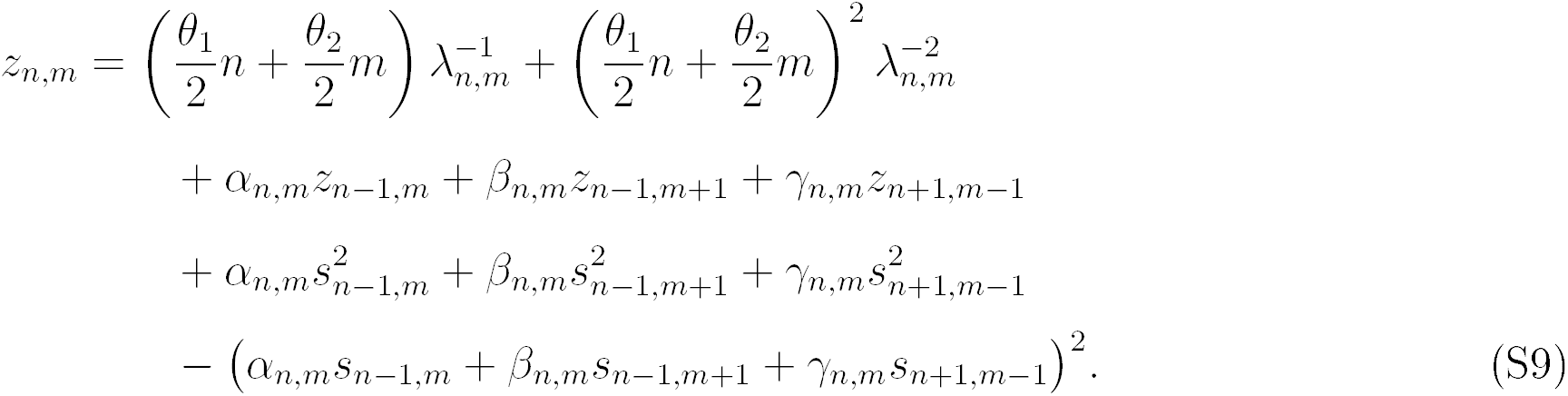

*Proof of Proposition S1.5.* Let *σ*_1_ denote the number of mutations that occur until time *τ*_1_, which was defined in the proof of Proposition S1.1. Given *τ*_1_ = *t,* we know that *σ*_1_ is the sum of two independent Poisson random variables with parameters *θ*_1_*nt* and *θ*_2_*mt,* respectively. As in the previous proof we obtain

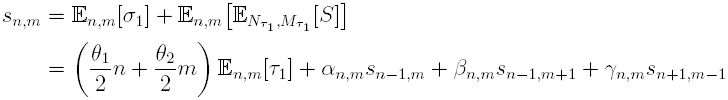

and

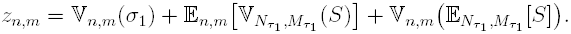

Once more using the law of total variance we obtain

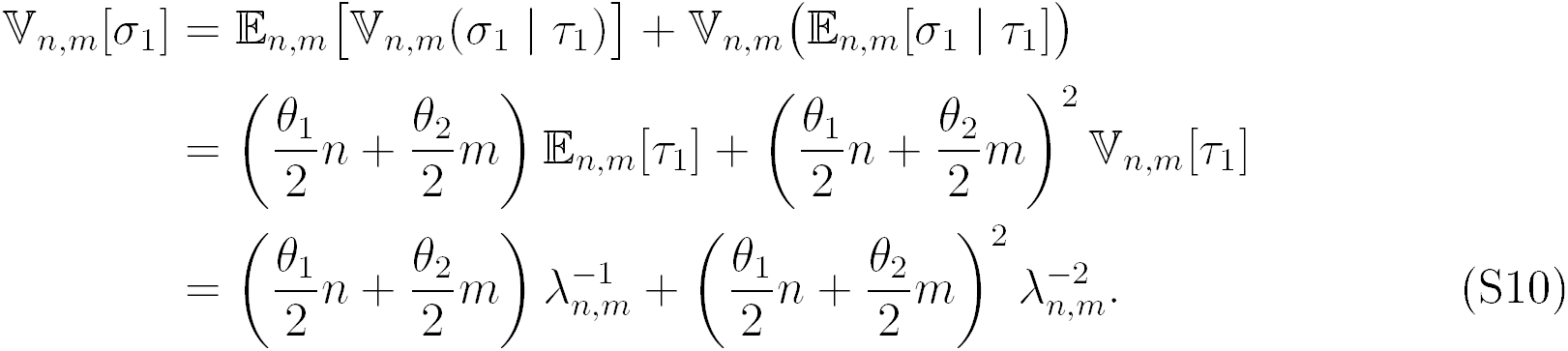

The same calculations as in the proof of Proposition S1.1 lead to

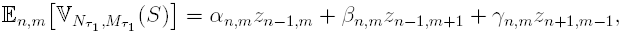

and

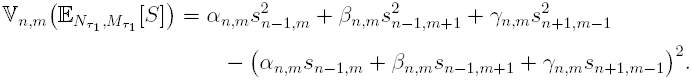

#### Expected value of average pairwise differences (*π*)

Recall the definition, given in (19), of *π*.

##### Proposition S1.6.

For *n, m ∊* ℕ _0_ *we have*

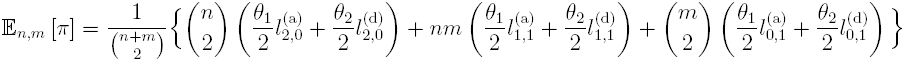

*Proof of Proposition S1.6.* By definition

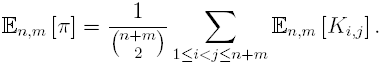

When compairing two individuals their *pairwise* differences in the infinite sites model coincide with the number of mutations that occured along the branches of their corresponding sub-tree and are thus given the product of the mutation rate and length of the branches. Therefore, 𝔼_*n*,*m*_[*K*_*i*,*j*_] actually only depends on whether *i, j* are dormant or active individuals. We obtain

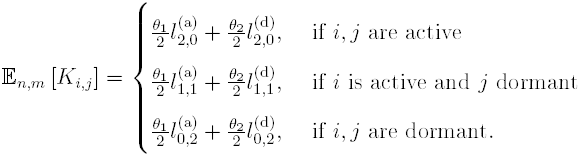

Substituting this into the above equation, the result follows.

#### S2 Solving the recursions nunerically

Since all the recursions have the same general form, a generic method for solving them numerically will now be given. The idea is to use standard linear algebra methods to solve the standard linear system 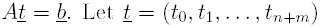 denote the vector of quantities we are solving for, where we order them according to number of active lines. For any given number *n* of active blocks and *m* of inactive blocks, so the current total number of blocks is *n* + *m*, write 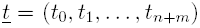 where *t*_*i*_ ≡ *t*_*i*,*n*+*m*-*i*_, and write *l*:= *n* + *m*. Let ***A***, ***B***, ***C*** denote square (*ℓ* + 1) × (*ℓ* + 1) matrices whose rows and columns are enumerated from 0; with non-zero terms *a*_*i*-*1*,*i*_ = *α_i,£-i_*, *b*_*i*,*i*-1_ = *β*_*i*_,*l*-*i*, *c*_*i*,*i*+1_ = *γ*_*i*,*l*-*i*_, and let ***I*** denote a (l + 1) × (*£* + 1) identity matrix. Assume, by way of example, we are solving the recursion (10) for expected time to most recent common ancestor. Define the vector 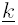 with elements *k*_*i*_ = 1*/λ_i,£-i_*, and 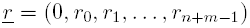 where *r*_*j*_ = *t*_*j*,*n*+*m*-*j*-1_.

The recursion in Proposition (S1.1) can now be written

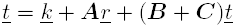

Assuming we solve for *t* iteratively, starting from *n*+*m* = 2, 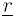 is a vector of known constants; hence

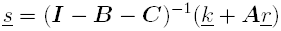

where ***I*** − ***B*** − ***C*** should be non-singular and (***I*** − ***B*** − ***C***)^−1^ easily computable.

Similar methods may be applied to the other recursions.

#### S3 Relative expected lengths of external branches

**Table SI.**
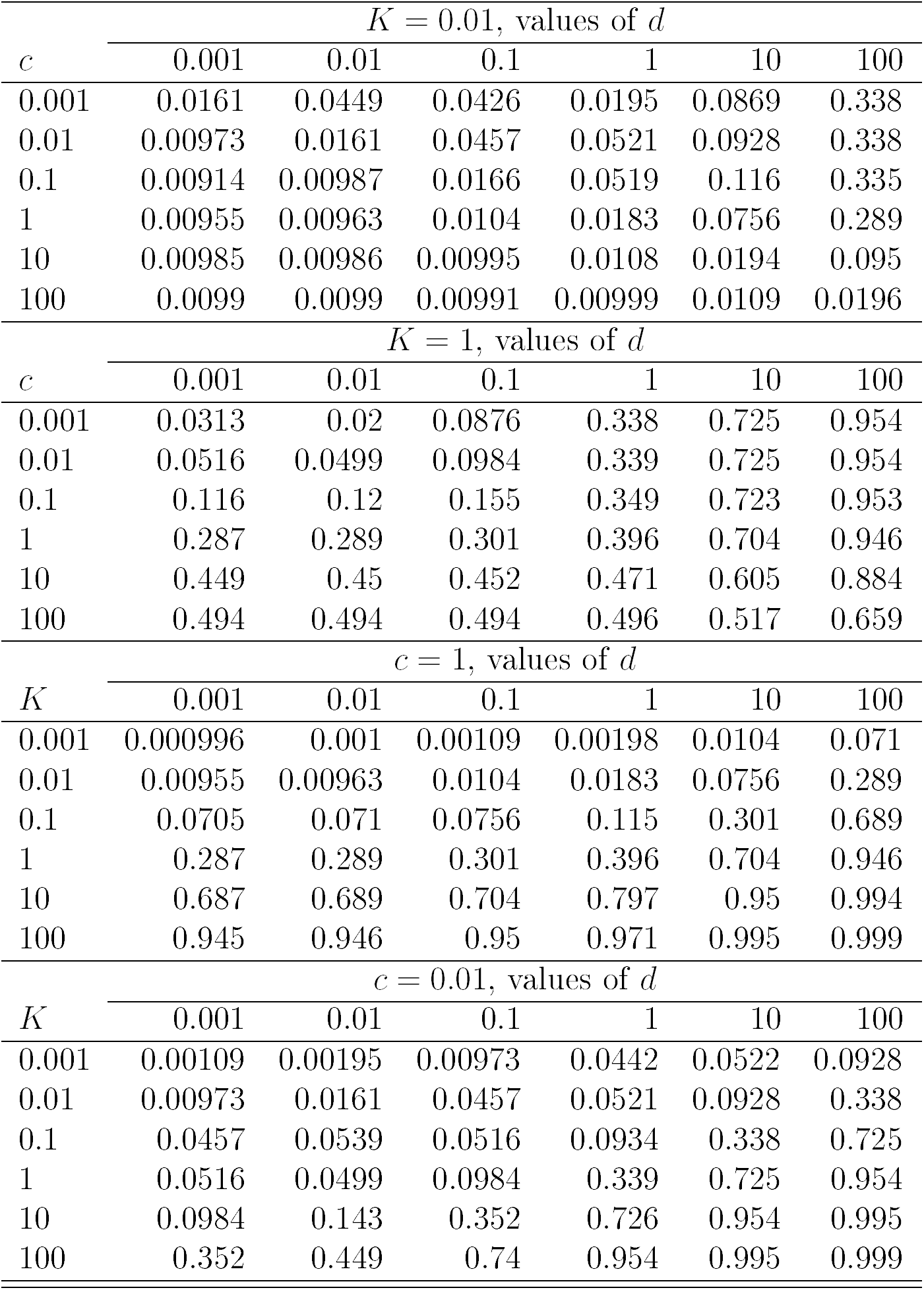
The relative expected lengths of external branches 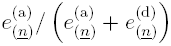 from Prop. S1.3 with sample configuration 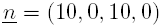.

#### S4 Expected length of external branches

**Table S2.** The expected total lengths of external branches 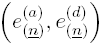 from Prop. S1.3 with sample configuration 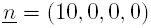, as a function of *c* and *K*. The expected length *e*_(*n*)_ = 2 when associated with the Kingman coalescent (Fu, 1995).

| $c = 1$, values of $d$ | | | |
| --- | --- | --- | --- |
| $K$ | 0.001 | 1 | 100 |
| 0.001 | (1.22e+03, 1.22e+06) | (610, 3.05e+05) | (14.2, 141) |
| 0.01 | (124, 1.24e+04) | (63, 3.15e+03) | (3.4, 3.36) |
| 0.1 | (14.3, 143) | (8.28, 41.4) | (2.18, 0.216) |
| 1 | (3.41, 3.4) | (2.77, 1.39) | (2.02, 0.02) |
| 10 | (2.18, 0.218) | (2.1, 0.105) | (2, 0.00198) |
| 100 | (2.02, 0.0202) | (2.01, 0.01) | (2, 0.000198) |
| $K = 0.01$, values of $d$ | | | |
| $c$ | 0.001 | 1 | 100 |
| 0.001 | (56.7, 2.83e+03) | (2.03, 0.203) | (2, 0.002) |
| 0.01 | (102, 9.28e+03) | (2.68, 2.65) | (2.01, 0.0201) |
| 0.1 | (111, 1.1e+04) | (11.5, 104) | (2.12, 0.211) |
| 1 | (124, 1.24e+04) | (63, 3.15e+03) | (3.4, 3.36) |
| 10 | (174, 1.74e+04) | (158, 1.44e+04) | (17.9, 163) |
| 100 | (198, 1.98e+04) | (196, 1.94e+04) | (100, 5.01e+03) |

#### S5 Expected nornalised branch lengths

**Figure S1:**
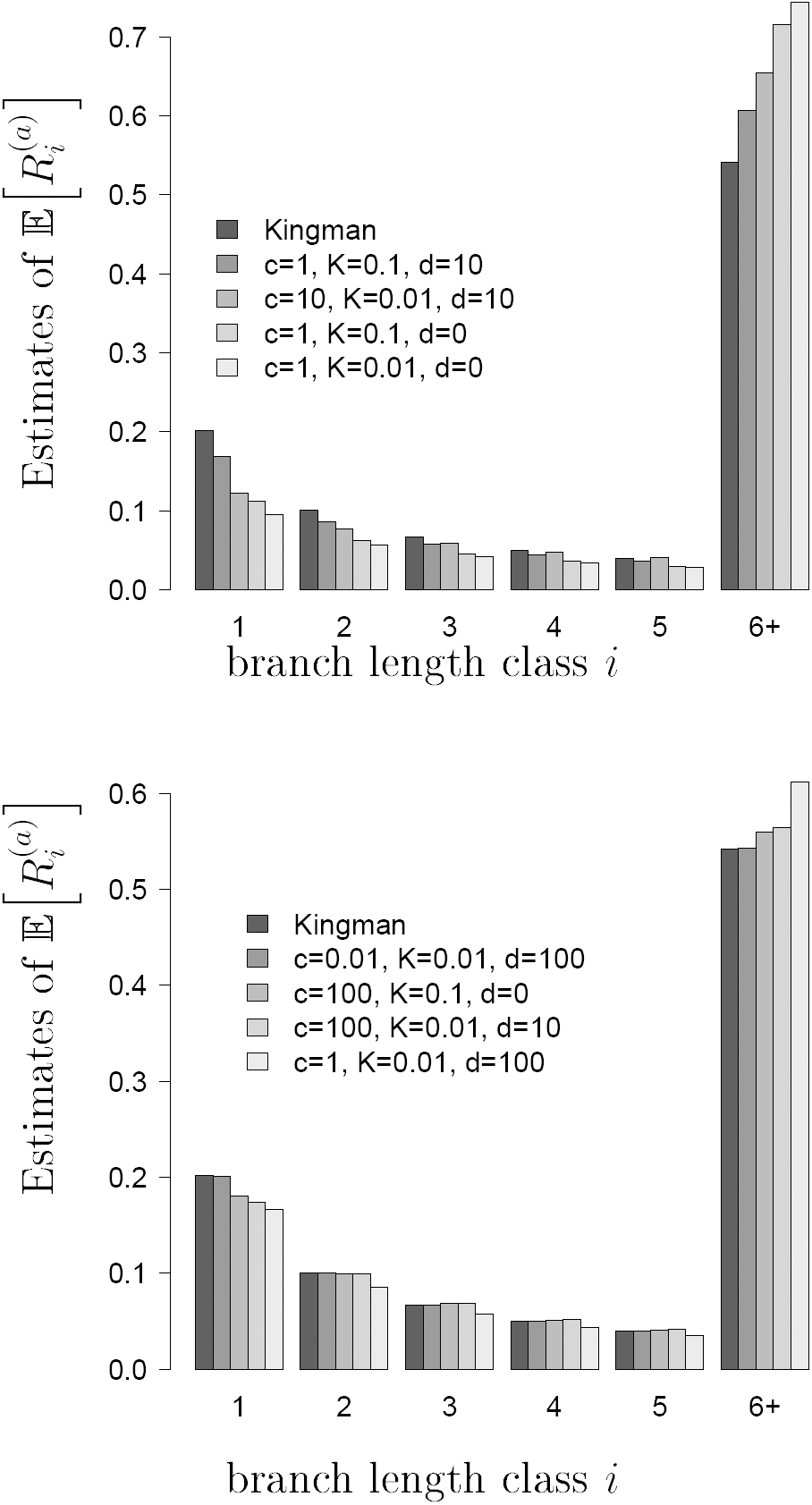
Estimates of the expected normalized branch lengths 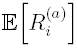 with 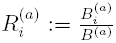 with 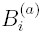 denoting the random total length of active branches subtending *i* leaves, and *B*^(a)^ the sum of *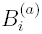*; with all *n* = 100 sampled lines assumed active, and values of *c*, *K*, *d* as shown. The values labelled 6+ denote the collected tail 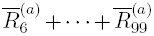. All estimates based on 10^5^ replicates.

#### S6 Expected normalised site-frequency spectrum

**Figure S2:**
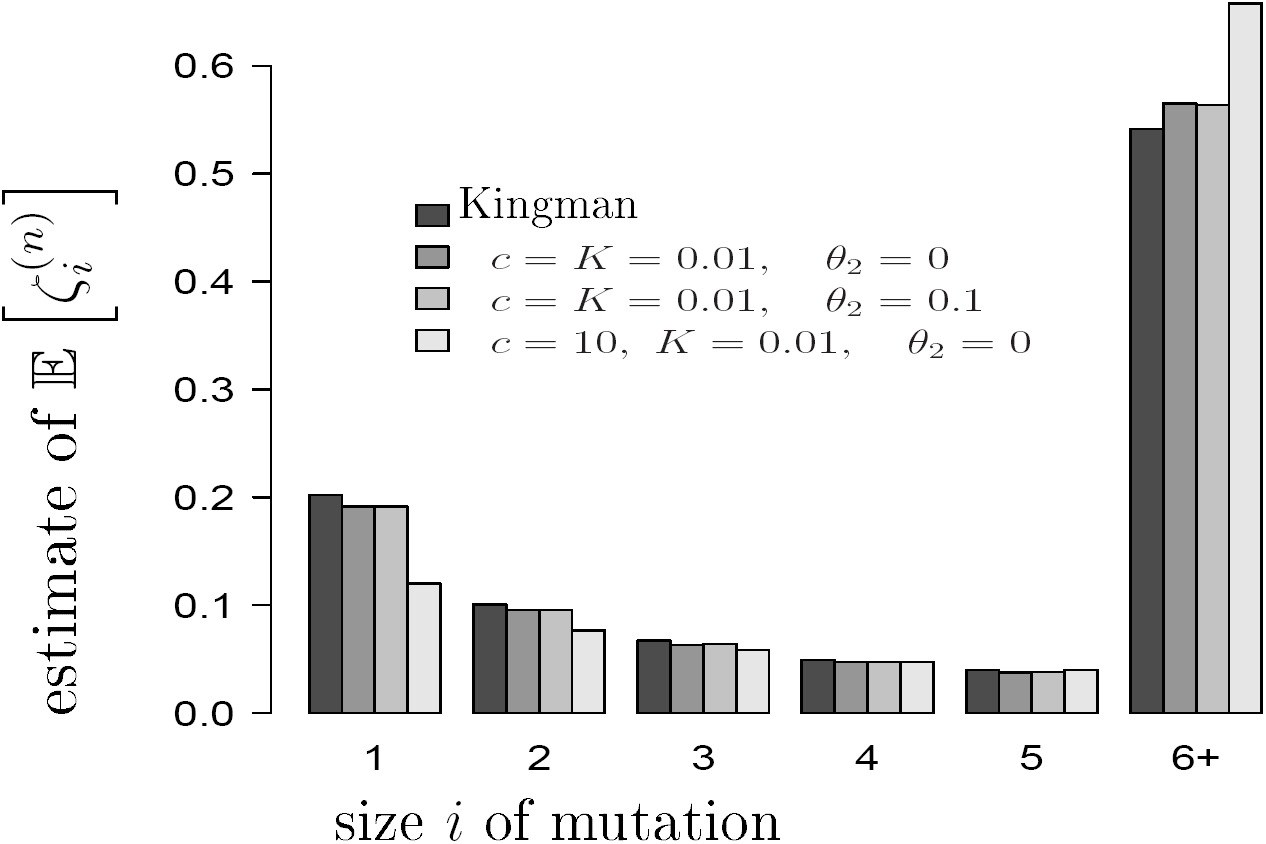
Estimates 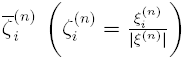 where 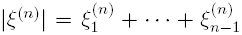 denotes the total number of segregating sites, of expected normalized spectra 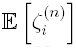 with all *n* = 100 sampled lines assumed active, active mutation rate *θ*_1_ = 2, and with *c*, *K*, and inactive mutation rate *θ _2_* as shown. The entries labelled ‘6+ ’ represent the collected tail 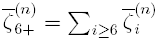. Estimates are based on 10^5^ replicates

#### S7 Tajina’s statistic *D*_T_ (24)

**Figure S3:**
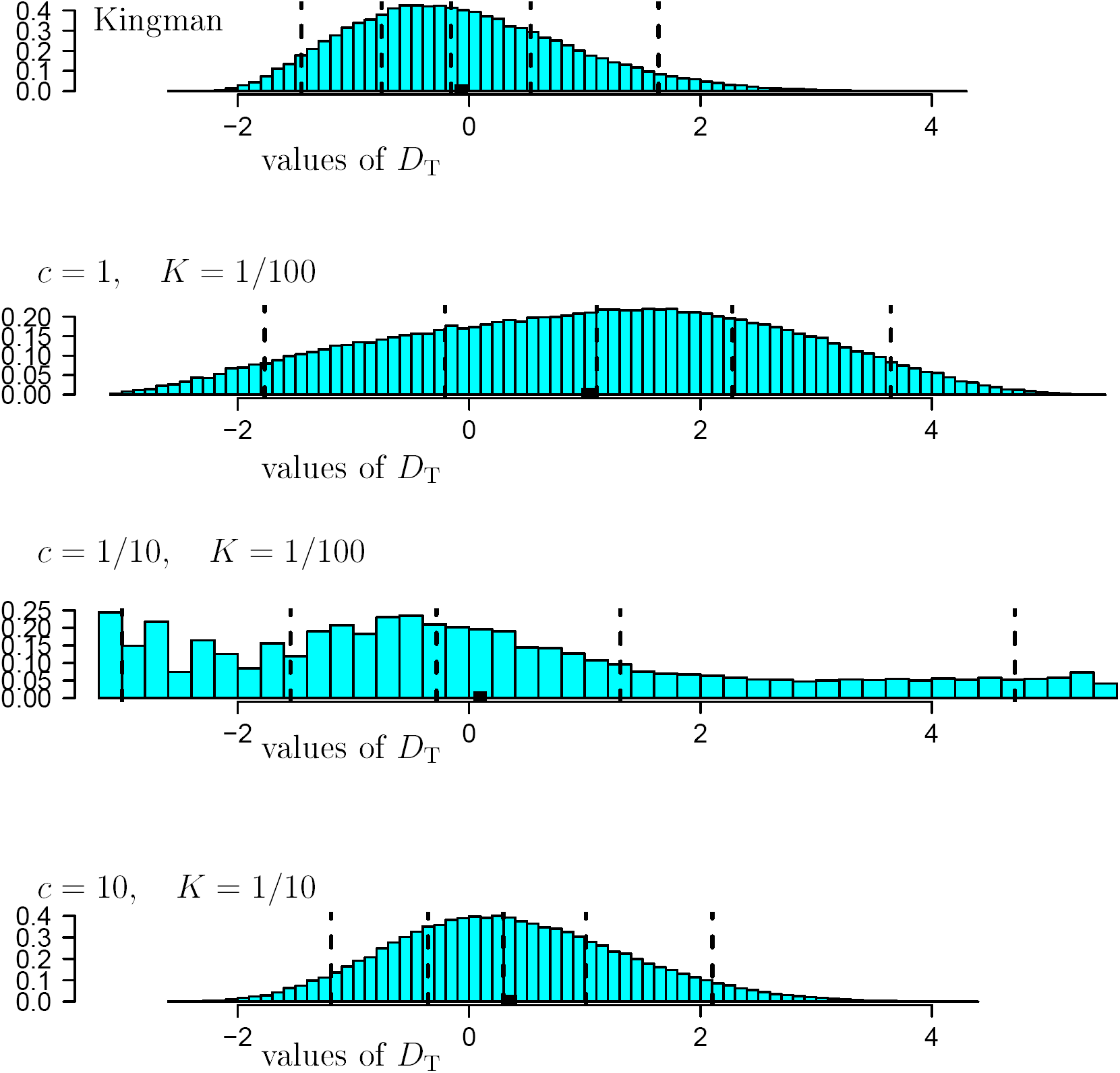
Estimates of the distribution of Tajima’s statistic *D*_T_ (24) with all *n* = 100 sampled lines assumed active, *θ*_1_ = 2, *θ*_2_ = 0. The vertical broken lines are the 5%, 25%, 50%, 75%, 95% quantiles and the black square (▮) denotes the mean. The entries are normalised to have unit mass 1. The histograms are drawn on the same horizontal scale. Based on 10^5^ replicates.

